# Layer-specific control of inhibition by NDNF interneurons

**DOI:** 10.1101/2024.04.29.591728

**Authors:** Laura Bella Naumann, Loreen Hertäg, Jennifer Müller, Johannes J. Letzkus, Henning Sprekeler

**Affiliations:** Institute of Science Technology Austria (ISTA), Klosterneuburg, Austria; Modelling of Cognitive Processes, Berlin Institute of Technology, Germany; Institute for Physiology, Faculty of Medicine, University of Freiburg, Germany; Spemann Graduate School of Biology and Medicine (SGBM), University of Freiburg, Germany; Faculty of Biology, University Freiburg, Germany; BrainLinks-BrainTools, Institute for Machine-Brain Interfacing Technology (IMBIT), University of Freiburg, Germany; Center for Basics in NeuroModulation (NeuroModul Basics), University of Freiburg, Germany; Bernstein Center for Computational Neuroscience, Berlin, Germany; Science of Intelligence, Research Cluster of Excellence, Berlin, Germany

## Abstract

Neuronal processing of external sensory input is shaped by internally-generated top-down information. In the neocortex, top-down projections predominantly target layer 1, which contains NDNF-expressing interneurons, nestled between the dendrites of pyramidal cells (PCs). Here, we propose that NDNF interneurons shape cortical computations by presynaptically inhibiting the outputs of somatostatin-expressing (SOM) interneurons via GABAergic volume transmission in layer 1. Whole-cell patch clamp recordings from genetically identified NDNF INs in layer 1 of the auditory cortex show that SOM-to-NDNF synapses are indeed modulated by ambient GABA. In a cortical microcircuit model, we then demonstrate that this mechanism can control inhibition in a layer-specific way and introduces a competition for dendritic inhibition between NDNF and SOM interneurons. This competition is mediated by a unique mutual inhibition motif between NDNF interneurons and the synaptic outputs of SOM interneurons, which can dynamically prioritise different inhibitory signals to the PC dendrite. NDNF interneurons can thereby control information flow in pyramidal cells by redistributing dendritic inhibition from fast to slow timescales and by gating different sources of dendritic inhibition, as exemplified in a predictive coding application. This work corroborates that NDNF interneurons are ideally suited to control information flow within cortical layer 1.

## Introduction

The neocortex receives a multitude of inputs that provide both sensory information and internally generated signals such as behavioural relevance (Sarter et al., 2005; Zhang et al., 2014) or expectations (Rao and Ballard, 1999; Friston, 2012). These different information streams need to be filtered and integrated to form accurate sensory perceptions and produce appropriate behavioural responses. While sensory inputs are typically relayed from the thalamus (“bottom-up”), inputs from other cortical and subcortical areas carry memory-or context-related signals (“top-down”; Gilbert and Sigman, 2007; Roth et al., 2016; Pardi et al., 2020) and mostly target the uppermost layer of cortex – layer 1 (L1). Cortical L1 stands apart from other layers for its absence of excitatory cell bodies, instead containing the dendrites of pyramidal cells located in deeper layers (Vogt, 1991; Felleman and Van Essen, 1991; Mitchell and Cauller, 2001; Larkum, 2013) and inhibitory interneurons (INs; Schuman et al., 2021). With the identification of the genetic marker NDNF (neuron-derived neurotrophic factor) that selectively labels L1 INs (Abs et al., 2018), one class of these cells have become accessible for specific characterisation and manipulation, but how NDNF INs contribute to cortical computation remains an open question.

Inhibitory INs differ in their morphology, electrophysiology, peptide expression and connectivity within the circuit. The most prevalent and well-studied IN types are parvalbumin-expressing (PV), somatostatin-expressing (SOM) and vasointestinal-peptide-expressing (VIP) INs. PV, SOM and VIP INs form a characteristic connectivity pattern within the cortical microcircuit that is remarkably similar across sensory cortex and species (Pfeffer et al., 2013; Campagnola et al., 2022; Schneider-Mizell et al., 2023). Their unique properties make them suitable for specialised functions (Isaacson and Scanziani, 2011; Kepecs and Fishell, 2014; Hangya et al., 2014; Fishell and Kepecs, 2020). For example, SOM INs exert a powerful inhibition to the PC dendrite, controlling the propagation of input signals to the soma (Markram et al., 2004). PV INs, on the other hand, inhibit the perisomatic region of PCs and are thus implicated in providing stability by balancing excitatory inputs (Ferguson and Gao, 2018). VIP INs inhibit SOM INs, thus disinhibiting PCs. Since VIP INs are driven by top-down inputs, this disinhibition has been linked to behavioural state modulation and plasticity (Pi et al., 2013; Hangya et al., 2014; Wilmes et al., 2016; Krabbe et al., 2019; Wilmes and Clopath, 2019).

Despite their strategic location among PC dendrites and top-down inputs, L1 INs have gained less attention compared to other inhibitory INs, largely owing to their sparse distribution and, until recently, the absence of a specific marker (Abs et al., 2018). NDNF INs in L1 receive top-down and neuromodulatory inputs in mouse and human neocortex (Abs et al., 2018; Poorthuis et al., 2018; Pardi et al., 2020) but unlike VIP INs provide slow inhibition to PC dendrites (Tamás et al., 2003; Schuman et al., 2019; Hartung et al., 2023). They can inhibit other INs but do not reciprocate the inhibition they receive from SOM INs (Abs et al., 2018; Hartung et al., 2023, Fig. 1A). Morphologically, NDNF INs are neurogliaform cells (Poorthuis et al., 2018; Abs et al., 2018; Schuman et al., 2019). A distinguishing feature of these cells is that they mediate GABAergic volume transmission, which besides causing slow postsynaptic effects can target presynaptic GABA_B_ receptors (Oláh et al., 2009; Pardi et al., 2020). Presynaptic inhibition via GABAergic volume transmission was recently identified as a mechanism to locally control inputs such as top-down projections to L1 (Pardi et al., 2020; Naumann et al., 2022).

**Figure 1.**
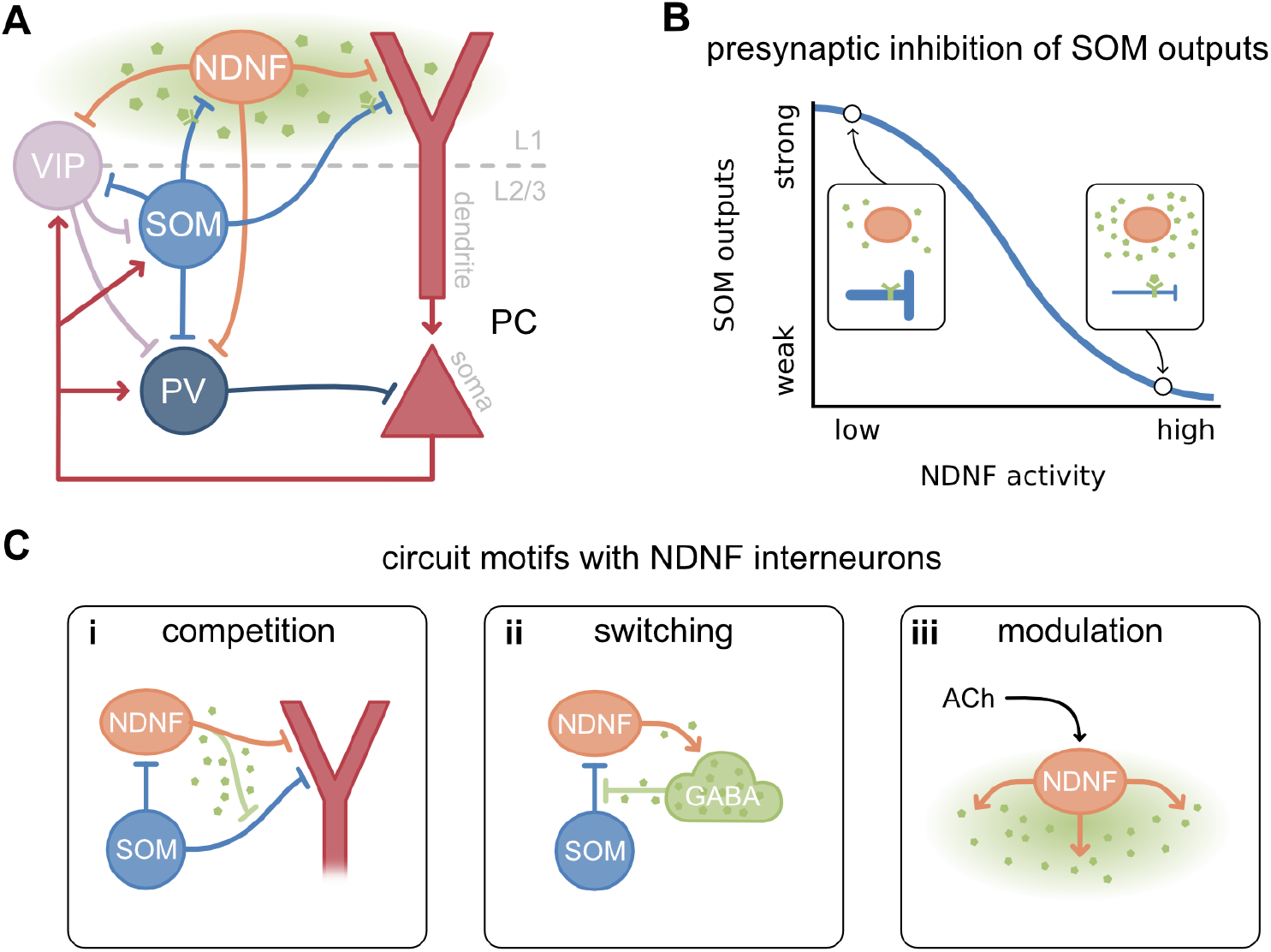
Microcircuit model with NDNF INs mediating GABAergic volume transmission. **A**. Schematic of the cortical microcircuit model with NDNF, SOM, PV and VIP INs and two-compartment PCs. **B**. Illustrative influence of NDNF activity on SOM outputs. Insets show low and high GABA release (i.e. NDNF activity) respectively. **C**. Circuit motifs with NDNF INs and presynaptic inhibition.

Motivated by these findings we hypothesised that NDNF INs shape cortical processing by exerting a presynaptic modulation of SOM IN outputs in L1 via GABAergic volume transmission (Fig. 1 A, B). Due to the location of NDNF IN output synapses (Abs et al., 2018), we propose that this modulation is limited to L1, implying that NDNF INs control the inhibition provided by SOM INs in a layer-specific way. To explore this hypothesis, we combine computational modelling and in vitro electrophysiology. First, we expand a model of the canonical cortical microcircuit by NDNF INs and GABAergic volume transmission. To validate our assumption that SOM outputs are presynaptically inhibited by ambient GABA, we performed whole-cell patch clamp recordings from genetically identified NDNF INs in mouse auditory cortex slices. Our experiments confirm that SOM synapses to NDNF INs in L1 are indeed modulated by presynaptic GABA_B_ receptors. Using our microcircuit model, we show that this mechanism introduces novel functional motifs (Fig. 1C): (i) Stimulating NDNF INs replaces SOM inhibition to PC dendrites with NDNF inhibition, creating a competition for the control of dendritic activity. (ii) NDNF INs locally counteract the inhibition they receive from SOM INs, a motif that can amplify signals to NDNF INs and function as a bistable switch between NDNF INs and SOM outputs. Since NDNF and SOM INs mediate inhibition on different timescales this redistributes inhibition in time. (iii) Neuromodulatory projections targeting NDNF INs can dynamically shape the signal processing in PCs. We show that modulating NDNF IN activity affects the relative balance of sensory (i.e. bottom-up) and top-down inputs to PCs, dynamically changing what PCs respond to in a predictive coding example.

## Results

Given the unique properties of NDNF INs and their strategic location among PC dendrites and top-down inputs in L1, we wondered how they contribute to cortical computation. To study how NDNF INs interact with the local circuit, we first introduced them into a classical cortical microcircuit model Pfeffer et al. (2013); Hertäg and Sprekeler (2019). The rate-based model contains a population of excitatory PCs and the four main IN types PV, SOM, VIP and NDNF (Fig. 1A). Each PC consists of two coupled compartments representing the soma and the dendrite, whereas INs consist of a single compartment (see Methods). Connection strengths and probabilities between the neuron types are motivated by electrophysiological studies of cortical layer 1-3 (Pfeffer et al., 2013; Campagnola et al., 2022, see Materials and Methods) and established microcircuit models (Litwin-Kumar et al., 2016; Yang et al., 2016; Hertäg and Sprekeler, 2019; Wilmes and Clopath, 2019). We accounted for GABAergic volume transmission from NDNF INs by modelling the GABA concentration in L1 (Fig. 1A, green cloud). The model assumes that the GABA concentration increases with NDNF IN activity and mediates slow inhibition of the PC’s dendrite. NDNF inhibition to PV and VIP INs is synaptic but weak, consistent with electrophysiological findings (Hartung et al., 2023). To model presynaptic inhibition of SOM outputs in L1, we include a release factor that multiplicatively scales the strength of SOM synapses and decreases with the GABA concentration (Fig. 1B, see Materials and Methods). We assume that GABAergic volume transmission is restricted to L1 (Oláh et al., 2009; Abs et al., 2018), affecting only the connections of SOM INs to the PC dendrite and NDNF INs, without impacting their connections to PV and VIP INs in lower layers, including layer 2-3.

### Experiments confirm the influence of NDNF INs on SOM outputs

Our main assumption is that SOM outputs in L1 are controlled by NDNF INs. In our model, SOM IN synapses to PC dendrites and to NDNF INs are presynaptically modulated by ambient GABA that is released by NDNF INs. A necessary prerequisite of the model is that the release probability of these synapses is modulated by presynaptic GABA receptors. The two main targets of SOM outputs in L1 are PC dendrites and NDNF INs. Synapses from SOM INs to PCs indeed express presynaptic GABA_B_ receptors in the hippocampus, and SOM-induced inhibitory currents in PCs are markedly reduced by the application of the GABA agonist baclofen (Booker et al., 2020). However, it is unknown whether synaptic transmission from SOM to NDNF INs in the auditory cortex is modulated by presynaptic GABA_B_ receptors.

To directly address this assumption of the model, we performed electrophysiological recordings in the auditory cortex *in vitro*. To this end, we crossed mice expressing Cre recombinase under the SOM promoter with a strain expressing Flp recombinase under the NDNF promoter (Abs et al., 2018). Stereotactic injection of adeno-associated viral vectors (AAVs) into the auditory cortex was employed to achieve SOM IN-specific expression of the optogenetic activator ChR2, and NDNF IN-specific expression of tdTomato, a fluorescent marker protein (Fig. 2A). This allowed us to perform whole-cell patch clamp recordings from genetically identified NDNF INs in layer 1 of the auditory cortex in acute brain slices, while at the same time enabling optical stimulation of SOM INs with millisecond precision (Fig. 2B).

**Figure 2.**
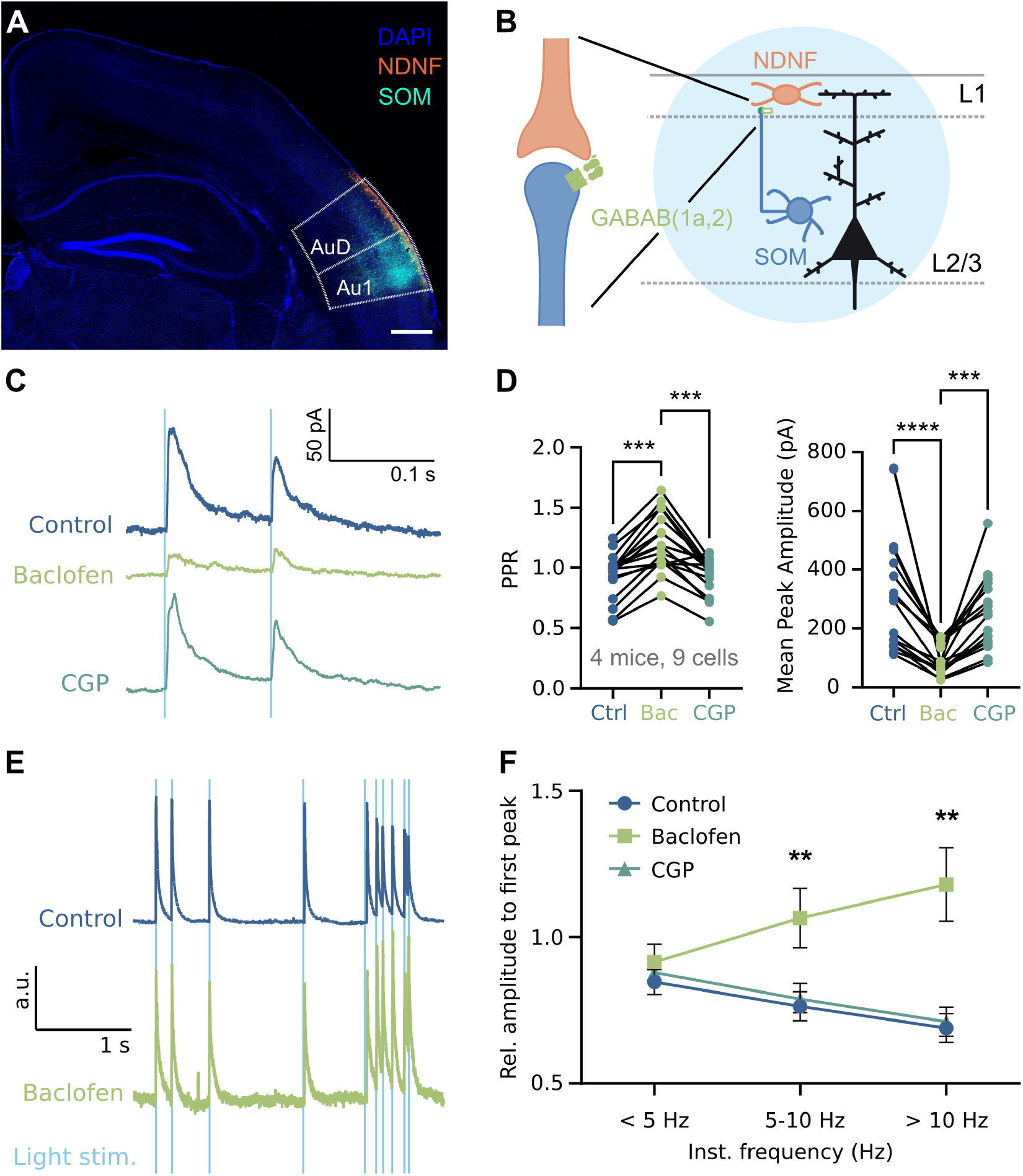
Experiments confirm the influence of NDNF INs on SOM outputs. **A**. Confocal microscope image showing expression of tdTomato in NDNF INs and ChR2-EYFP in SOM INs. Scale bar 500*μ*m. **B**. Hypothesis: NDNF INs in L1 modulate SOM IN inputs through presynaptic GABA_B_ receptor-mediated inhibition. Light blue circle represents optogenetic full field stimulation. **C**. Representative IPSCs during paired pulse stimulation at 10 Hz (0.5ms pulse). **D**. Paired pulse ratio and mean peak amplitude for Control (ACSF), Baclofen and CGP55845. **E**. Representative IPSCs during naturalistic stimulation, normalised to first response. **F**. Normalised response amplitude, grouped according to instantaneous frequency. Data shown as averages of 10 sweeps for (C), (D), and (E). Data shown as mean ± SEM. * *p <* 0.05, ** *p <* 0.01, *** *p <* 0.001, **** *p <* 0.0001.

Optogenetic activation (0.5ms pulses) of SOM INs caused robust inhibitory postsynaptic currents (IPSCs) in almost all NDNF INs tested (91%), consistent with the observed strong connectivity between these IN types (Abs et al., 2018). We minimised possible postsynaptic effects of GABA_B_ receptor activation by using Cesium-based intracellular solution (Pardi et al., 2020, Supp. Fig. S1). Bath application of the selective GABA_B_ receptor agonist Baclofen (10 *μ*mol/l) strongly reduced the amplitudes of IPSCs (Fig. 2C, D right, mean Ctrl 312.9 pA, Baclofen 95.06 pA, CGP 248.1 pA; Ctrl vs Baclofen *p <* 0.0001, Ctrl vs CGP *p* = 0.3681, Baclofen vs CGP *p* = 0.0009; Friedman test with Dunn’s multiple comparisons test). Paired-pulse stimulation at 10 and 20 Hz further revealed an increase in paired-pulse ratio (PPR) under GABA_B_ receptor activation (Fig. 2C, D left, mean Ctrl 0.9331, Baclofen 1.212, CGP 0.9267; Ctrl vs Baclofen *p* = 0.0001, Baclofen vs CGP *p* = 0.0001, Control vs CGP *p >* 0.9999; Friedman Test with Dunn’s multiple comparisons test), consistent with presynaptic effects. Since both stimulation frequencies showed comparable effects, these data were pooled (see Supp. Fig. S1 for individual plots). Moreover, both the effects on IPSC amplitude and PPR were completely reversed by the selective GABA_B_ receptor antagonist CGP55845 (3 *μ*mol/l). Both the decrease in IPSC amplitude and the increase in PPR suggest the presence of presynaptic GABA_B_ receptors on synaptic terminals of SOM INs that target NDNF INs. In particular, the PPR is the most widely-used metric to quantify changes in presynaptic release probability (Regehr, 2012; Tsodyks and Markram, 1997). Therefore, these data demonstrate that GABA_B_ receptors dynamically and powerfully control the release probability at synaptic contacts from SOM INs to NDNF INs.

Importantly, presynaptic control can not only dynamically reconfigure the strength of a connection, but also its frequency transfer function (Tsodyks and Markram, 1997). We therefore investigated how presynaptic GABA_B_ receptors control transmission under more naturalistic conditions. To this end, we used a spike train that was previously recorded in vivo (Pardi et al., 2020) to define a naturalistic stimulation protocol comprising ten different instantaneous frequencies (Fig. 2E, ranging from 1 Hz to 26.77 Hz). The naturalistic stimulation revealed that pharmacological activation of presynaptic GABA_B_ receptors indeed shifts the maximum of the frequency transfer function between SOM INs and NDNF INs from low (*<* 5 Hz) under control and GABA_B_ receptor antagonism to high during baclofen application (Ctrl vs Baclofen *p* = 0.0098 for 5 − 10 Hz, Ctrl vs Baclofen *p* = 0.0038 for *>* 10 Hz, RM Two-way ANOVA with Tukey’s multiple comparisons test). Together, these data demonstrate that presynaptic GABA_B_ receptors robustly and dynamically control both the strength and the frequency transfer function at SOM IN connections to NDNF INs. Our experiments on SOM-to-NDNF synapses together with earlier work on SOM-to-PC synapses (Booker et al., 2020) suggest that SOM outputs in L1 are indeed under the control of NDNF INs through GABAergic volume transmission. In the following, we investigate how this added level of computational flexibility at SOM synapses affects circuit function.

### Competition between SOM-and NDNF-mediated dendritic inhibition

Having established that NDNF IN can modulate SOM outputs in L1 via GABAergic volume transmission, we asked how this mechanism affects the cortical microcircuit at the functional level. The two primary targets of SOM outputs in L1 are PC dendrites and NDNF INs. First, we focus on the role of modulating SOM outputs to PC dendrites, which are also inhibited by NDNF INs (Fig. 3A, top). As a result, increasing the activity of NDNF INs reduces the inhibition from SOM INs to pyramidal cell dendrites, replacing it with NDNF-mediated dendritic inhibition (Fig. 3A, middle graph). In other words, NDNF INs and SOM outputs compete for dendritic inhibition. Notably, SOM INs and their outputs in lower layers are not directly affected (Fig. 3A, top graph), provided the effect of GABAergic volume transmission is restricted to cortical L1. Indirect effects on SOM INs can nevertheless occur due to recurrent interactions within the microcircuit, either through disinhibition via VIP INs (NDNF-VIP-SOM pathway) or changes in PC activity (NDNF-PC-SOM pathway, cf. Fig. 1A, Supp. Fig. S6).

**Figure 3.**
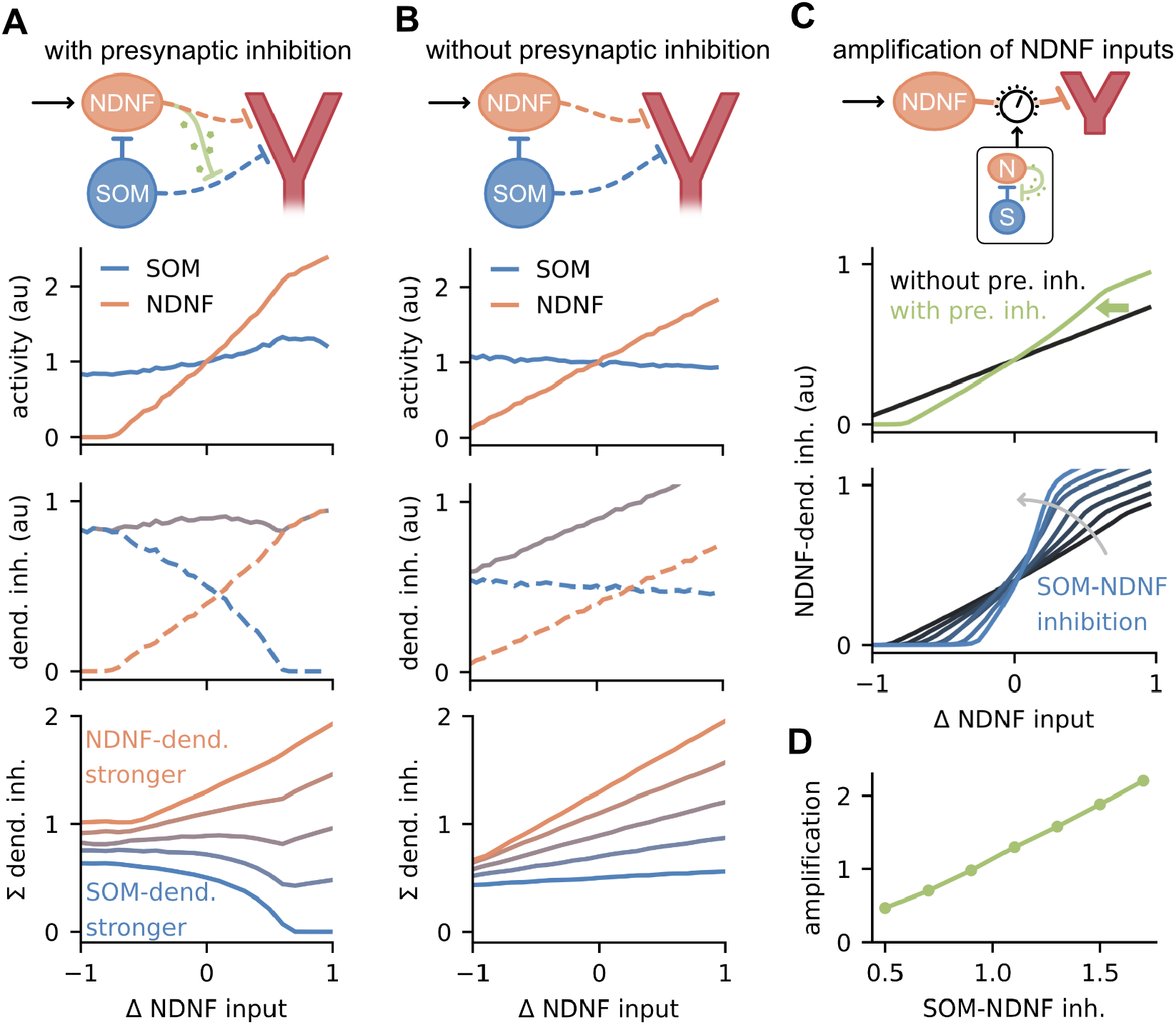
Competition between SOM-and NDNF-mediated dendritic inhibition. **A./B**. Model behaviour with and without presynaptic inhibition for different levels of NDNF input relative to baseline. Top: Sub-circuit consisting of SOM INs, NDNF INs, and PC dendrite. Second row: Activity of NDNF and SOM INs. Third row: Dendritic inhibition from SOMs (blue), NDNFs (orange), and both combined (grey). Bottom: Total dendritic inhibition for varying strengths of NDNF-to-dendrite inhibition. More orange colours indicate a stronger NDNF-to-dendrite synaptic weight and blue colours indicate a weaker weight. **C**. Top: Illustration of the amplification of NDNF input by the NDNF-SOM motif. Middle: NDNF-dendrite inhibition with/without presynaptic inhibition as a function of NDNF input. Bottom: Same as above but for varying strengths of SOM-NDNF inhibition (*w*_NS_ between 0.5 and 1.7). **D**. Amplification of NDNF input as a function of SOM-NDNF inhibition. Amplification is quantified as the log ratio between the NDNF input-output slope with and without presynaptic inhibition (shown in C).

Whether the total dendritic inhibition is higher when NDNF or SOM inhibition dominates depends on their relative strength (Fig. 3A, bottom). When SOM-to-dendrite inhibition is stronger, stimulation of NDNF INs scales down SOM outputs and replaces them with weaker NDNF-to-dendrite inhibition, thereby decreasing the overall dendritic inhibition. Conversely, when NDNF-to-dendrite inhibition is stronger, stimulation of NDNF INs increases the overall dendritic inhibition. The competition for dendritic inhibition stems from presynaptic inhibition on SOM synapses to PC dendrites as NDNF INs do not directly inhibit SOM INs. Without presynaptic inhibition, stimulating NDNF INs does not modulate the SOM-to-dendrite inhibition, thus only increasing the overall dendritic inhibition (Fig. 3B). Monitoring dendritic activity in response to NDNF stimulation can therefore serve as an indicator of the strength of presynaptic inhibition on SOM outputs and the relative strength of SOM-compared to NDNF-mediated dendritic inhibition. Activation of NDNF INs decreases dendritic inhibition only if presynaptic inhibition and SOM-to-dendrite inhibition are sufficiently strong.

At first glance, NDNF INs seem to be at a disadvantage when competing for dendritic inhibition, because they are unidirectionally inhibited by SOM INs (Abs et al., 2018). However, our experiments revealed that SOM-to-NDNF synapses can also be modulated by NDNF-mediated presynaptic inhibition. This provides NDNF INs with an intriguing mechanism to counteract the inhibition they receive from SOM INs by effectively scaling it down (Fig. 3C, top). From a mathematical point of view, SOM-to-NDNF outputs and GABAergic volume transmission via NDNF INs form an unconventional and layer-specific “mutual inhibition” motif: SOM outputs inhibit NDNF INs and in return, NDNF INs presynaptically inhibit SOM outputs via GABAergic volume transmission (see Materials and Methods). Although this is not a classical mutual inhibition motif between inhibitory populations, we found that it displays similar properties. Depending on the strength of the mutual inhibition, this motif can amplify small differences in the input and become bistable (Hertäg and Sprekeler, 2019). Indeed, we find that presynaptic inhibition amplifies the NDNF-to-dendrite inhibition evoked by stimulating NDNF INs (Fig. 3C, top & middle). The amplification increases with the SOM-to-NDNF inhibition (Fig. 3C, bottom & D), which – together with presynaptic inhibition – determines the strength of the mutual inhibition. Our model suggests that modulation of SOM-to-PC synapses by NDNF INs can gradually control the balance of SOM-and NDNF-mediated dendritic inhibition, introducing an effective competition between the two pathways that is restricted to L1. At the same time, the NDNF-mediated dendritic inhibition is amplified by an unconventional form of mutual inhibition between NDNF INs and SOM outputs.

### NDNF INs can act as a switch for dendritic inhibition

We wondered if the NDNF-SOM motif could be pushed to bistability similar to classical mutual inhibition circuits (Hertäg and Sprekeler, 2019). To test this, we provided transient input pulses to NDNF INs and observed their effect on the circuit (Fig. 4A). If the SOM-to-NDNF inhibition is sufficiently strong, positive input pulses lead to long-lasting increases and negative pulses to long-lasting decreases in NDNF activity (Fig. 4B-D). Yet, for weak SOM-to-NDNF inhibition, transient inputs to NDNF INs do not have lasting effects (Fig. 4E), regardless of the pulse strength (Fig. 4C). Stimulation of NDNF INs (e.g. positive pulses) causes an increase in their activity and thus the ambient GABA concentration. As a result, presynaptic inhibition scales down SOM outputs, reducing the SOM-mediated inhibition to NDNF INs. The relief from SOM IN-mediated inhibition allows for further increases in NDNF IN activity. Transient inputs can therefore switch NDNF INs to an active or inactive state. Note that the SOM IN activity is not affected by the switching (Fig. 4D, top), unlike in a classical mutual inhibition motif. Instead, the NDNF-SOM circuit exhibits winner-take-all behaviour between NDNF IN activity and SOM *outputs*. As SOM outputs to the PC dendrites are also modulated by NDNF-mediated presynaptic inhibition, this “mutual inhibition” motif can serve as a switch for the source of dendritic inhibition (Fig. 4D, bottom).

**Figure 4.**
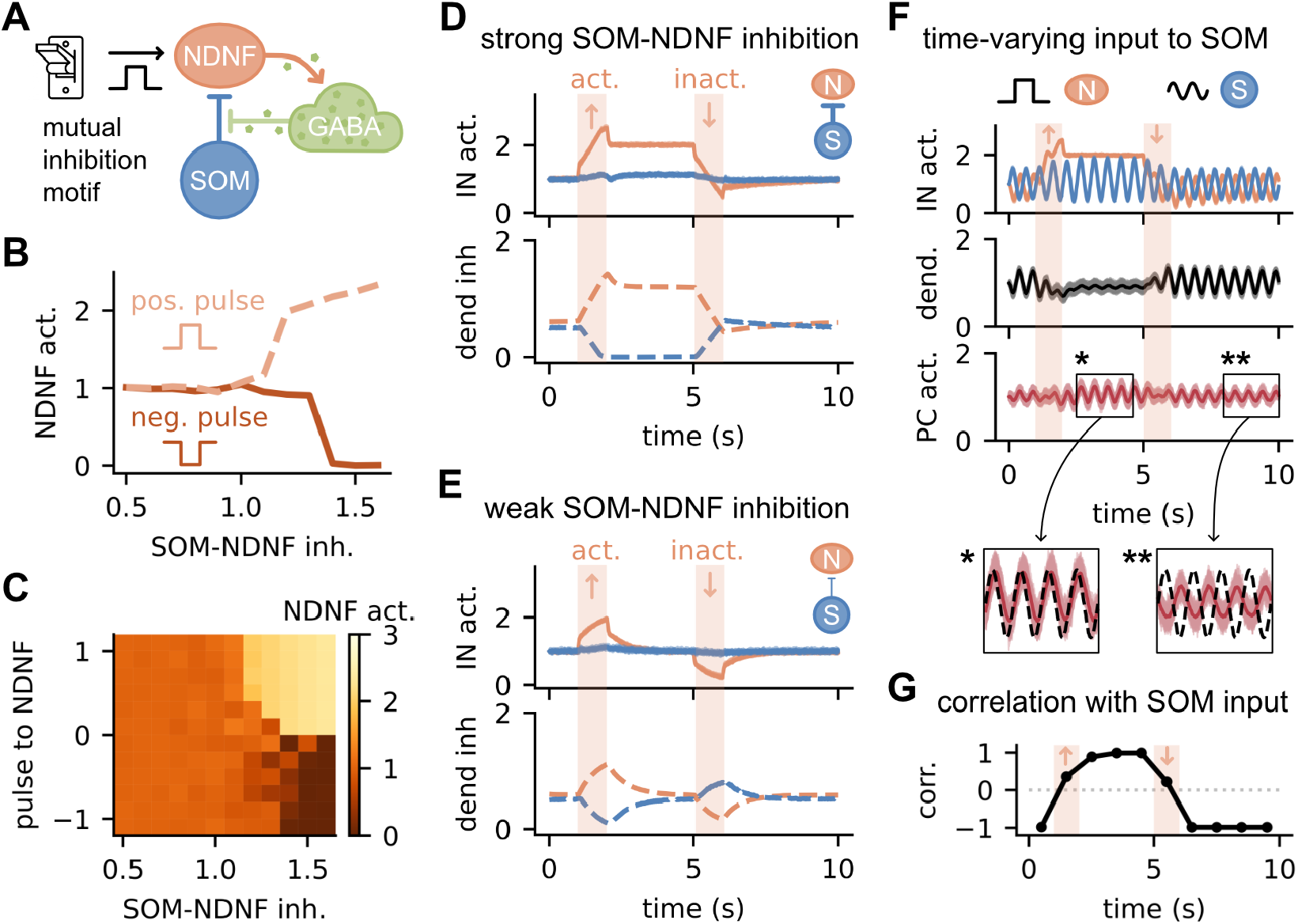
NDNF INs can act as a switch for dendritic inhibition. **A**. Schematic illustration of the switch between NDNF INs and SOM outputs. **B**. Steady-state NDNF activity as a function of SOM-NDNF inhibition strength after a positive (dashed) or negative (solid) pulse to NDNFs. **C**. Same as B but for different pulse strengths and signs. **D**. Time course of SOM and NDNF activity (top) and the dendritic inhibition they exert (bottom) when NDNFs are switched on and off. SOM-NDNF inhibition is strong (*w*_NS_ = 1.2). **E**. Same as (D) but for weak SOM-NDNF inhibition (*w*_NS_ = 0.7). **F**. SOM and NDNF activity (top), dendritic inhibition (center) and PC activity (bottom) in response to time-varying input to SOMs and pulses to NDNFs. **G**. Correlation of PC activity with SOM input from (F).

From a functional perspective, why should it matter whether PC dendrites are inhibited by NDNF or SOM INs? NDNF and SOM INs display similar output connectivity patterns within the circuit, inhibiting PC dendrites, VIP and PV INs. However, they receive different inputs and thus represent different signals: While SOM INs receive bottom-up sensory input (Urban-Ciecko and Barth, 2016), NDNF INs are targeted by top-down feedback inputs (Abs et al., 2018; Pardi et al., 2020). To illustrate how switching between NDNF-and SOM-mediated dendritic inhibition can influence signal transmission in PCs, we provided a time-varying signal (i.e. a sine wave) to SOM INs and repeated our switching experiment (Fig. 4F). When NDNF INs are switched to a more active state by a positive input pulse, the sine signal in the dendrite is markedly attenuated, because SOM outputs to NDNF INs and PC dendrites are inhibited presynaptically (Fig. 4F, centre). Switching NDNF INs back to the baseline recovers the signal in the dendrites. Remarkably, the time-varying signal provided to SOM INs still appears in the PC activity, even when SOM-to-dendrite synapses are inhibited. The reason is that the inhibition from SOM to PV INs in lower layers is not affected by presynaptic inhibition and thus SOM INs influence the PC somata through the SOM-PV-PC pathway (cf. Fig. 1A). In contrast to direct SOM-mediated inhibition, this pathway disinhibits the PCs, such that the sign of the time-varying input signal to SOM INs is reversed by the switch. Conversely, when SOM inhibition dominates the dendrites, the PC signal is inverted compared to the SOM input (Fig. 4F, bottom insets). We quantified the signal inversion by computing the correlation between SOM input and PC response, which flips from negative to positive when NDNF INs are switched on (Fig. 4G).

Collectively, these results demonstrate that the SOM-NDNF IN motif can be pushed to form a bistable switch for dendritic inhibition that dynamically changes the signals represented in PCs in response to transient inputs. This mechanism is particularly compelling when NDNF and SOM INs transmit different information to PCs such as bottom-up or top-down signals.

### Redistribution of dendritic inhibition in time

Besides receiving distinct input signals, NDNF and SOM INs differ in their temporal dynamics, in particular in the timescale of inhibition to PC dendrites. While SOM INs provide direct synaptic inhibition mediated by GABA(A) receptors, NDNF INs tend to inhibit PC dendrites via GABAergic volume transmission targeting both GABA(A) and extrasynaptic GABA_B_ receptors (Tamás et al., 2003; Oláh et al., 2009; Abs et al., 2018; Schuman et al., 2019). Consequently, the postsynaptic currents elicited by NDNF INs show slower dynamics. In our model, this difference is captured by the GABA concentration that slowly increases with NDNF IN activity and mediates the inhibition to the dendrite (Fig. 5A, B). The slow NDNF IN-mediated inhibition takes time to build up, which – in combination with other inhibitory pathways in the circuit – results in a multi-phased response in the PCs (Fig. 5C). The faster GABA(A)-mediated NDNF-PV-PC pathway causes a brief initial increase in PC activity, followed either by a further increase or a decrease depending on the strength of NDNF-to-dendrite inhibition (Supp. Fig. S4& S4). The termination of the stimulus can evoke another PC response phase due to the interplay of slow NDNF-dendrite inhibition, presynaptic inhibition of SOM outputs and fast NDNF-PV-PC disinhibition (Supp. Fig. S6).

**Figure 5.**
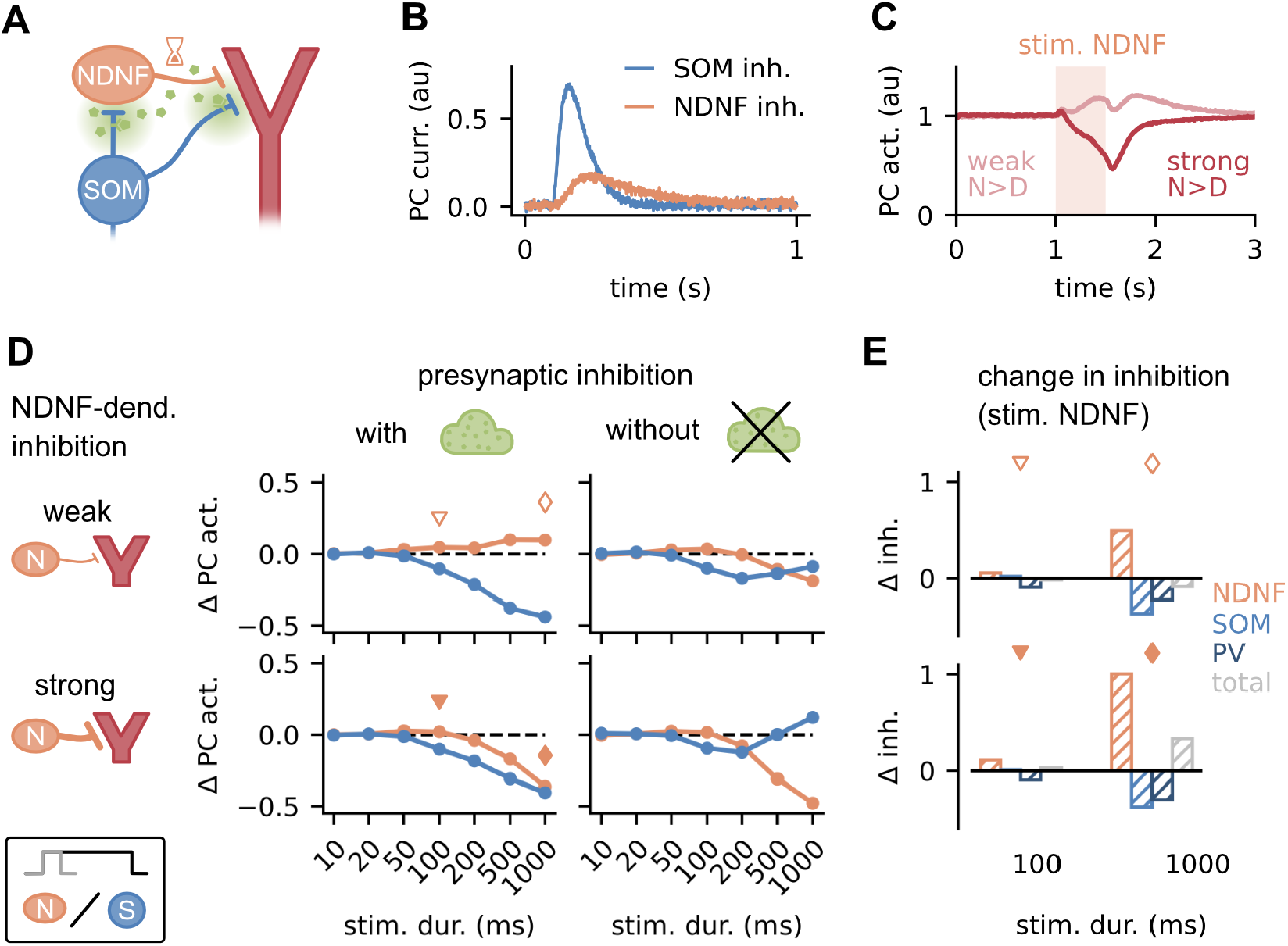
Redistribution of dendritic inhibition in time. **A**. Sub-circuit consisting of NDNF INs, SOM INs, GABAergic volume transmission and PC dendrites. **B**. Inhibitory current in PCs in response to SOM and NDNF IN stimulation. **C**. Mean PC activity in response to NDNF IN stimulation (orange box) for weak and strong NDNF-to-dendrite inhibition. **D**. Response of PCs to constant NDNF and SOM IN stimulation of different durations. The same experiment is shown for weak/strong NDNF-dendrite inhibition and with/without presynaptic inhibition. **E**. Contribution of inhibition provided to PCs by NDNFs, SOMs, PVs and total inhibition when stimulating NDNF INs in (D), measured by the change compared to baseline. Triangles and diamonds denote corresponding data points from (D).

To systematically study the downstream effects of NDNF compared to SOM stimulation, we provided pulses of varying lengths to NDNF or SOM interneurons. We find that the effect of these pulses depends both on their length and the circuit configuration. First, we focus on the circuit responses in our model with presynaptic inhibition (Fig. 5D, left). Stimulation of SOM INs generally decreases the PC activity, with longer stimulation evoking larger decreases (Fig. 5D, left). The PC response to NDNF IN stimulation is more complex and varies with the strength of the NDNF-to-dendrite inhibition (weak or strong compared to other weights in the circuit, see Methods). For weak NDNF-to-dendrite inhibition, PC activity increases with longer NDNF IN stimulation (Fig. 5D, top left). The underlying reason is that activating NDNF INs primarily disinhibits PCs, via two routes. Firstly, NDNF INs reduce SOM-to-dendrite inhibition via presynaptic inhibition. Secondly, they disinhibit PC somata via the NDNF-PV-PC pathway (Fig. 5E, top, diamond; Supp. Fig. S6). The importance of these pathways depends on the circuit parameters. When NDNF-to-dendrite inhibition is strong, this direct inhibitory contribution dominates the disinhibitory pathways (Fig. 5E, bottom), thus decreasing PC activity for longer NDNF IN stimulation. However, short stimuli can still cause a weak increase in the PC response (Fig. 5E, bottom, triangle), because the two pathways operate on different timescales. The direct synaptic NDNF-to-PV inhibition (i.e. PC disinhibition) is faster than the GABAergic volume transmission from NDNF INs that inhibits PC dendrites and SOM outputs in L1.

Our model predicts that stimulating NDNF INs with varying stimulus durations can be used to determine the relative strength of NDNF-to-dendrite inhibition. Long input stimuli should have opposite effects on PC activity for weak compared to strong NDNF-to-dendrite inhibition (cf. Fig. 5D left, diamonds; Supp. Fig. S4& S5). Similarly, the PC responses to IN stimulation can be used to identify the presence or contribution of presynaptic inhibition in the circuit: Without presynaptic inhibition, NDNF INs predominantly inhibit PCs, because they do not counteract the SOM-mediated dendritic inhibition (Fig. 5D, right). Furthermore, NDNF INs cannot counteract the inhibition from SOM INs (cf. Fig. 4) such that stimulating SOM INs reduces NDNF IN activity and their inhibition of the dendrite. This implies that SOM IN stimulation counterintuitively increases the PC response when NDNF-to-dendrite inhibition is strong.

The model shows that stimulating NDNF and SOM INs can have diverse downstream effects depending on the relative balance of multiple inhibitory and disinhibitory pathways in the microcircuit. The stimulus duration plays a crucial role because NDNF INs mediate inhibition on longer timescales (Tamás et al., 2003; Schuman et al., 2019; Hartung et al., 2023). The predictions from our model could be tested in future experiments by stimulating NDNF and SOM INs and using the PC responses as a unique signature to delineate relative pathway strengths and the contribution of presynaptic inhibition in the microcircuit.

### NDNF INs enable switching between prediction-responsive and mismatch neurons

We have shown that NDNF INs can control inhibitory pathways in L1 through GABAergic volume transmission and thereby modulate signal transmission to PCs. To illustrate how this layer-specific control affects cortical processing at a computational level, we turned to a predictive coding example. The idea of predictive coding is that the brain aims to predict sensory information using internally generated predictions (Bell, 1981; Rao and Ballard, 1999; Friston, 2012; Keller and Mrsic-Flogel, 2018). Deviations from predicted signals cause prediction errors that can be used to refine the inner model of the world and therefore improve future predictions. Prediction error (i.e. mismatch) responses have been widely observed (Schultz and Dickinson, 2000; Keller and Mrsic-Flogel, 2018). For example, a subset of PCs in layer 2/3 of rodent primary visual cortex specifically responds to mismatches between observed visual flow and the expected flow from motor commands (Keller et al., 2012; Attinger et al., 2017). Similar responses were found in auditory cortex (Eliades and Wang, 2008; Keller and Hahnloser, 2009). Recent theoretical work established how cortical microcircuits can give rise to mismatch responses, identifying the important role of multiple IN types to balance different sensory and prediction inputs (Hertäg and Sprekeler, 2020; Hertäg and Clopath, 2022). However, these models did not consider NDNF INs.

Motivated by our findings, we hypothesised that NDNF INs can dynamically modulate the responses in prediction error circuits. Because NDNF INs are driven by feedback and neuromodulatory inputs (including cholinergic inputs; Pardi et al., 2020; Hartung et al., 2023), they could shape prediction error responses depending on context or behavioural state. To test this idea, we tuned our cortical microcircuit model such that PCs respond to prediction errors, extending the previous prediction error circuit to include NDNF INs (Fig. 6A). As in previous work, we assume that sensory input projects to PC somata, PV and SOM INs. Conversely, top-down predictions cause inputs to PC dendrites and VIP INs.

**Figure 6.**
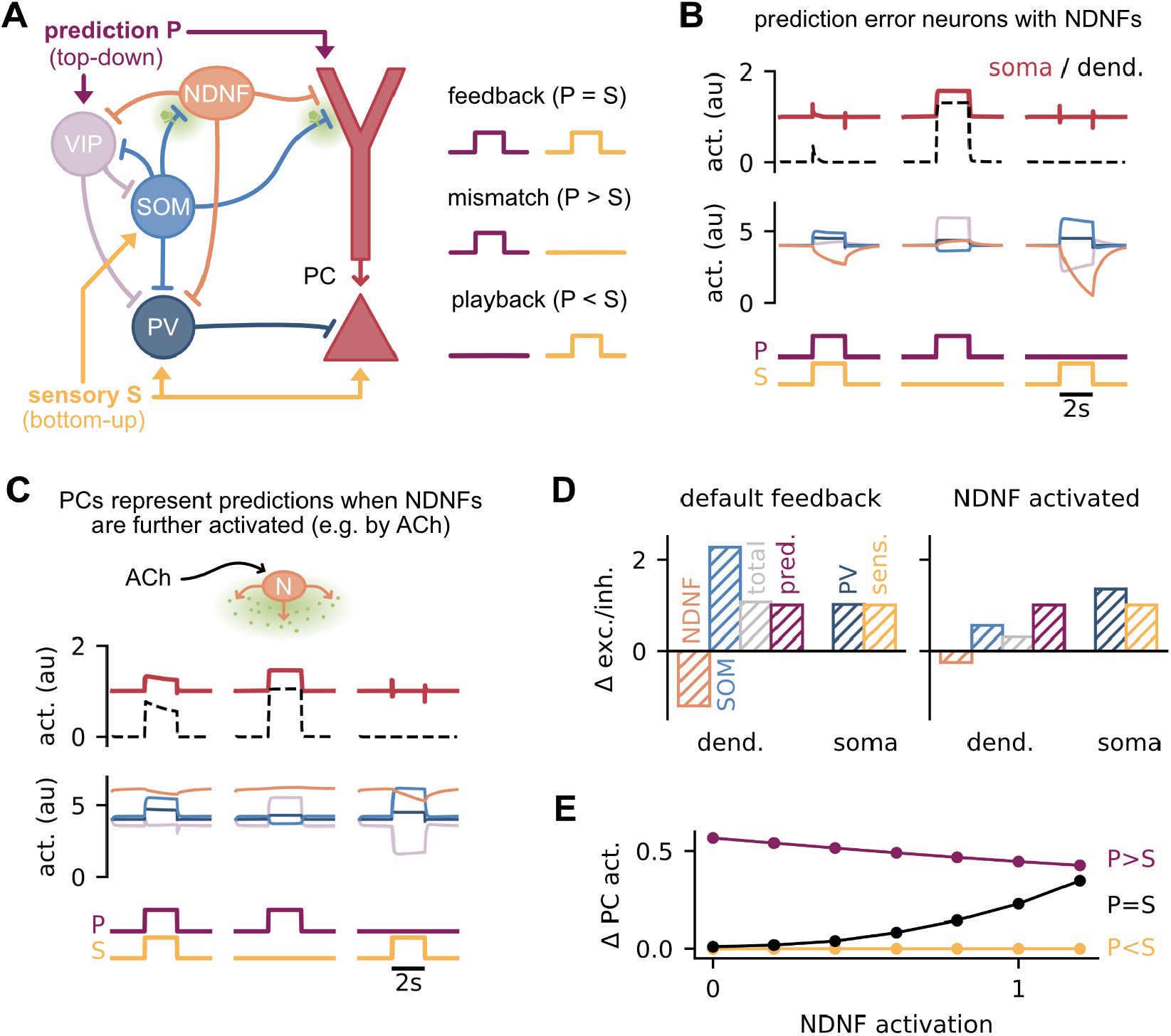
NDNF INs enable switching between prediction-responsive and prediction error neurons. **A**. Prediction error circuit with NDNF INs (left) and different conditions for predictive and sensory input (right). **B**. Responses of PC dendrites and somata (top) and four IN groups (center) to different input configurations. Colours correspond to the schematic in (A). **C**. Same as (B) but with NDNF INs activated, e.g. by cholinergic input. **D**. Change in excitatory and inhibitory inputs to dendrite and soma during the feedback phase in the default condition (left) and with NDNF INs activated (right). Colours correspond to (A). **E**. Change in PC activity as a function of NDNF activation for the three phases shown in (A).

We probed the responses of PCs to three different input combinations (see Fig. 6A, right). In the “feedback” condition, top-down input accurately predicts sensory inputs. In the “mismatch” condition, there is a top-down prediction but no sensory input. Finally, in the “playback” condition, sensory input is present but not the associated top-down prediction. Mismatch neurons should respond only when the prediction outweighs the sensory input (mismatch), but not when the sensory input is predicted or there is no prediction at all (feedback and playback; Hertäg and Sprekeler, 2020).

The PC responses in our predictive coding circuit with NDNF INs are consistent with those of mismatch neurons (Fig. 6B, top; Attinger et al., 2017; Hertäg and Sprekeler, 2020). PCs do not respond to conjunctive sensory and prediction input (feedback condition, Fig. 6B, left), because it is balanced out by the inhibitory pathways in the prediction error circuit: the total dendritic inhibition (SOM and NDNF-mediated) balances the prediction input at PC dendrites and the PV inhibition balances the sensory input at the PC somata (Fig. 6D, left). Sensory input alone (playback condition) does not evoke a PC response, because SOM Ins maintain inhibition to PC dendrites and PV INs counteract the sensory input at the soma (Fig. 6B, right). Yet, in the mismatch condition, the prediction input activates VIP INs, which disinhibits PC dendrites and hence leads to a mismatch response (Fig. 6B, centre).

What happens to the mismatch responses when the activity of NDNF INs is modulated, for instance, by cholinergic inputs to L1 (Poorthuis et al., 2018)? Since NDNF INs control the inhibition of SOM INs to the dendrite, we expect them to influence the predictions arriving at the dendrites of PCs. We found that activation of NDNF INs causes PCs to respond in the feedback condition (Fig. 6C, top). At first glance, this response is not intuitive since SOM INs still increase their activity due to the sensory input (cf. Fig. 6B and C). However, SOM-and NDNF-mediated dendritic inhibition is smaller compared to the control condition, because SOM outputs are inhibited by the NDNF INs (Fig. 6D, right). As a result, the total dendritic inhibition is outweighed by the prediction input, allowing PC dendrites to become active. PCs therefore respond to predictions regardless of sensory information instead of prediction errors. This behaviour critically depends on the modulation of SOM IN outputs. Without presynaptic inhibition, activating NDNF INs does not qualitatively change the mismatch responses (Supp. Fig. S7).

In summary, these findings demonstrate that NDNF INs can dynamically shape the prediction error circuit. Whether PCs respond to mismatches or predictions can depend on the level of NDNF IN activity (Fig. 6B, C). Gradually varying the input to NDNF INs enables a smooth transition between prediction and mismatch responses (Fig. 6E). NDNF INs receive feedback and cholinergic inputs that signal, for instance, arousal state (Letzkus et al., 2011; Brombas et al., 2014; Poorthuis et al., 2018; Malina et al., 2021). We conjecture that the dynamic modulation of predictive coding responses can hence have behavioural relevance. During low arousal states (i.e. baseline NDNF activity), the circuit represents prediction mismatches, alerting the animal of deviations from its predicted sensory input. However, during high arousal (i.e. elevated NDNF activity), the circuit assigns more relevance to internal signals such as predicted stimuli, providing a mechanism for recognising expected stimuli more rapidly (Mazzucato et al., 2019).

## Discussion

We showed that NDNF INs can modulate cortical information processing by controlling the outputs of SOM INs in L1. NDNF INs release ambient GABA, which targets presynaptic GABA_B_ receptors and thereby inhibits synaptic transmission (Pardi et al., 2020). We validated experimentally that this mechanism affects the synapses from SOM to NDNF INs by performing optogenetic stimulation, patch-clamp recordings and pharmacological manipulations in slices of mouse auditory cortex. Together with evidence of presynaptic inhibition of SOM-to-PC synapses (Booker et al., 2020), these findings support our hypothesis that SOM outputs in L1 are controlled by presynaptic inhibition. In a cortical microcircuit model that includes NDNF INs, we explored the effects of presynaptic inhibition of SOM outputs on the circuit dynamics and function. We found that NDNF INs can control inhibition in a layer-specific way, by targeting SOM outputs. NDNF INs also form a competitive circuit motif with SOM INs, in which NDNF INs counteract the unidirectional inhibition from SOM INs by presynaptically inhibiting the SOM inputs they receive. The motif can amplify small signals and form a bistable switch that enables shifting between NDNF-and SOM-mediated inhibition to the dendrite, thus dynamically changing the signal processing in PCs. Stimulation of NDNF INs can have diverse downstream effects depending on the temporal dynamics of the stimulation and relative connection strengths. Finally, we illustrated the functional relevance of NDNF INs in a predictive coding example. Modulating the activity of NDNF INs, e.g. by cholinergic inputs, shapes the representation of predictions and prediction errors in the circuit. Our results demonstrate that by controlling SOM inhibition in a layer-specific way, NDNF INs increase the functional flexibility of cortical circuits.

### Layer-specific control: plausibility and functional implications

In our model, NDNF INs have a layer-specific inhibitory effect. The main assumption is that presynaptic inhibition via GABAergic volume transmission affects SOM synapses within L1, an inhibition that selectively targets SOM outputs rather than SOM neurons themselves. This specificity is particularly intriguing for Martinotti cells that project an axon to L1 while their somata lie in deeper layers (L2/3 or L5 Urban-Ciecko and Barth, 2016; Tremblay et al., 2016). If GABAergic volume transmission is confined to cortical L1, the activity and thus the outputs of these cells in lower layers remain unaffected. This includes their projections to PV INs, which are thought to be essential to maintain an excitation/inhibition balance at the soma of PCs (Ferguson and Gao, 2018). Similarly, the layer-specificity of the mechanism enables controlling inputs to PC dendrites separately from the soma. As the soma typically receives bottom-up sensory inputs while the dendrite receives top-down contextual and behavioural input (Larkum et al., 2004; Gilbert and Sigman, 2007; Roth et al., 2016; Schuman et al., 2021), this layer-specificity provides a nuanced control over cortical information processing relevant for cognitive functions such as predictive coding (Fig. 6).

But how specific is GABAergic volume transmission? We conjectured that ambient GABA released by NDNF INs only acts within L1, the resident layer of NDNF INs. To mediate presynaptic inhibition, GABA released by NDNF INs must reach presynaptic GABA_B_ receptors at, for instance, SOM output synapses. One challenge for this diffusive form of signalling is uptake mechanisms that actively remove neurotransmitters around synapses and release sites (Isaacson, 2000). Thus, the amount of released neurotransmitters must be sufficient to overcome reuptake and diffuse to nearby synapses. The diffusion of GABA is further limited by physical obstacles, L1 being densely packed with dendritic and axonal arbours (Schuman et al., 2021). NDNF INs (morphologically neurogliaform cells) are particularly well suited to drive GABAergic volume transmission, because they have a high density of GABA release sites that are often not associated with a synapse (Overstreet-Wadiche and McBain, 2015). Moreover, a single action potential can cause large slow postsynaptic inhibitory currents (Oláh et al., 2009; Jiang et al., 2015; Tremblay et al., 2016). As the NDNF IN axons as well as their output synapses are largely constrained to L1 (Abs et al., 2018; Schuman et al., 2019), and GABA diffusion is limited physically as well as by reuptake mechanisms (Oláh et al., 2009), it is unlikely for GABAergic volume transmission to exert a meaningful effect below L1.

While VIP interneurons are predominantly located in layer 2-3 (Tremblay et al., 2016), they can reach lower L1 (Schuman et al., 2019). Hence, GABAergic volume transmission may affect the synapses of SOM to VIP INs at the border of L1 if they express presynaptic GABA_B_ receptors. Another potential target of presynaptic inhibition in L1 is the synapses from NDNF INs to the dendrite (Oláh et al., 2009). We found that the competition and bistability between SOM and NDNF INs is robust to GABAergic volume transmission targeting SOM-to-VIP or NDNF-to-dendrite synapses (Supp. Fig. S2& S3). Therefore, our results do not critically rely on modulating only a specific subset of synapses within L1.

In our model, the ambient GABA concentration and its effect on presynaptic release probability is homogeneous within L1. This assumption was motivated by the observation that NDNF IN axonal arbours extend over large horizontal distances in L1 (Jiang et al., 2015; Overstreet-Wadiche and McBain, 2015; Schuman et al., 2019) and that their activity tends to be correlated (Malina et al., 2021), suggesting that – despite the sparseness of NDNF Ins – ambient GABA release is relatively uniform across the cortical microcircuit. We did not model the spatial distribution of cells within the cortical circuit beyond their home layer (L1 or L2/3). Future work could explore the role of spatially heterogeneous GABAergic volume transmission in a model with spatial structure.

### NDNF and SOM INs: competing master regulators

In the model, NDNF INs exert a powerful and unique control over the cortical microcircuit. Our work supports the notion that NDNF INs serve as “master regulators” of the cortical column (Hartung et al., 2023), a role that has also been ascribed to Martinotti-type SOM INs (Jiang et al., 2015). Both NDNF and SOM INs inhibit many other cells in the circuit, yet they form different connectivity patterns with different output mechanisms (presynaptic or postsynaptic) and timescales (Jiang et al., 2015; Abs et al., 2018; Pardi et al., 2020), suggesting that they operate in different ways (Jiang et al., 2015). These properties can result in distinct downstream circuit effects (Fig. 5). Furthermore, SOM INs tend to receive local recurrent and feedforward inputs that contain sensory information (Urban-Ciecko and Barth, 2016), whereas NDNF INs are targeted by top-down feedback inputs that carry contextual, behavioural-state or memory-related signals (Abs et al., 2018; Pardi et al., 2020; Hartung et al., 2023). Although NDNF INs can respond to sensory stimulation (auditory and visual), their sensory responses are sensitive to previous experience and behavioural states (Abs et al., 2018; Malina et al., 2021). SOM and NDNF INs thus regulate local circuitry based on distinct signals. Notably, we showed that these two “master regulators” may interact via a unique mutual inhibition motif, creating a dynamic interplay that flexibly regulates the cortical microcircuit depending on behavioural states, for instance. Understanding how this interplay influences cortical signal processing to guide behaviour will require future theoretical work and experimental studies *in vivo*.

### Limitations of our theoretical and experimental approach

We focussed on the key features of the cortical circuit model and made several simplifying design choices. Inhibitory INs were modelled as single-compartment rate neurons, describing the activity of each neuron by its firing rate. Therefore, the model does not consider the timing of spikes or electrophysiological differences between the INs. NDNF INs tend to show a late-spiking behaviour (Schuman et al., 2019; Hartung et al., 2023), distinct from the low-threshold and adaptive spiking of SOM INs or the fast-spiking of PV INs (Tremblay et al., 2016). Including the electrophysiological properties of the different INs would add another layer of complexity to the circuit model. While our results should still hold in a spiking circuit model, we expect the timing of input spikes to play a larger role. In particular, the late spiking of NDNF INs could introduce an additional temporal filter, adding to their slow inhibitory action.

We modelled L1 cells as a homogeneous class of INs that express NDNF. While their characteristics are still an active area of research, several lines of evidence point to at least two electrophysiologically distinct classes of L1 NDNF INs (Schuman et al., 2019; Gouwens et al., 2020; Hartung et al., 2023). However, the exact delineation and whether subclasses can be identified by genetic markers (such as NPY) remains controversial and may depend on the brain area, model species and developmental stages (Hartung et al., 2023). Future theoretical work could investigate how NDNF INs from different subclasses or from a spectrum of properties influence the circuit dynamics.

To validate our central assumption regarding the impact of GABAergic volume transmission on SOM outputs, we conducted experiments in mouse auditory cortex slices. Employing the GABA_B_ receptor agonist baclofen to emulate the release of ambient GABA, we observed a marked reduction in SOM-to-NDNF synaptic transmission. The activation of presynaptic GABA_B_ receptors was confirmed by decreased IPSC amplitudes and an increased paired-pulse ratio. This substantiates our hypothesis that SOM synapses onto NDNF INs express presynaptic GABA_B_ receptors, inhibiting synaptic transmission upon activation. However, the experiment does not directly show that endogenous GABA released by local NDNF INs has the same effect. An experimental approach where a single NDNF IN is recorded whilst multiple surrounding NDNF INs are stimulated for GABA release, and SOM Ins are stimulated for investigating synaptic connectivity, would require spatially resolved dual channel optogenetic stimulation. This is beyond our current technical scope. Even if possible, the experimental approach would additionally be restricted by artificial *in vitro* conditions where GABA diffusion and uptake are altered by physiological conditions. Given our primary focus on investigating the functional consequences of NDNF and SOM IN interactions, we concentrated on verifying our core hypothesis – the modulation of SOM outputs by ambient GABA.

Similarly, our approach emphasised achieving a qualitative alignment between the model and experimental results, rather than a quantitative match. Quantitatively fitting the model to observed data would require tuning numerous unknown parameters whose biological equivalent is challenging to measure, such as the amount of GABA released by NDNF INs, its diffusion range and the time it takes to reach nearby GABA_B_ receptors or undergo reuptake. The challenge in measuring these biological equivalents underscores the need for innovative techniques. Recent advances in optogenetic targeting of neuromodulators and neurotransmitters, such as GABA, offer promising avenues for more accessible quantification (Marvin et al., 2019; Sabatini and Tian, 2020; Wu et al., 2022). Optogenetic manipulations could be extended to genetically engineered G-protein-coupled receptors like GABA_B_, mimicking their activation through endogenous GABA. Apart from serving as a new tool for optogenetic inhibition, such manipulations hold the potential to directly explore the effects of presynaptic inhibition, including SOM outputs, in behaving animals.

## Materials and Methods

To study how NDNF interneurons control inhibition in L1, we took a combined experimental and theoretical approach. We used computational modelling to frame our hypothesis that NDNF INs presynaptically inhibit the outputs of SOM INs in L1 and explore its functional implications. In addition, we tested our core hypothesis by performing electrophysiological recordings in slices from mouse auditory cortex in combination with optogenetic and pharmacological manipulations.

### Interneuron microcircuit model

Inhibitory neurons (NDNF, SOM, PV and VIP interneurons) are point neurons described by a single activity value, whereas excitatory PCs consist of two compartments representing the dendrite and the soma. The activity of each neuron/compartment evolves according to a rectified, linear differential equation describing how the different cell types influence each other akin to

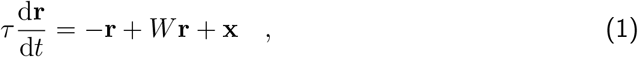

where r is the activity vector of all neurons, *τ* the time constant of the process, **x** the external input and *W* the synaptic interaction strength between neurons in the circuit. All neurons in the microcircuit are randomly connected with connection strengths and connection probabilities that are consistent with electrophysiological recordings (Pfeffer et al., 2013; Campagnola et al., 2022). In addition, neurons receive constant, external background input that ensures non-zero baseline activity.

To model the effect of NDNF IN-mediated GABAergic volume transmission, we introduce the ambient GABA concentration *c*_G_. The GABA concentration increases with the activity of all NDNF INs 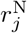 in the circuit with a time constant *τ*_G_:

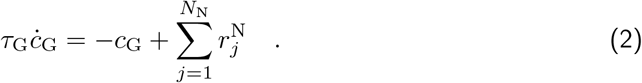

We include presynaptic inhibition by decreasing a release factor *p* with increasing GABA concentration, thereby capturing the inhibition of synaptic transmission via presynaptic GABA_B_. *p* evolves according to

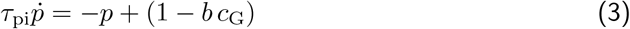

and is clipped between 0 and 1. The parameter *b* describes the strength of presynaptic inhibition and *τ*_pi_ its timescale. We scale the influence of SOM INs on both the PC dendrites and the NDNF INs by the release factor *p* such that the output synapses of SOM INs in L1 are modulated by presynaptic inhibition (see Fig. 1A).

Detailed descriptions of the model, the simulation experiments as well as model parameters can be found in the Supplementary Information.

### Experimental procedure

All mouse lines used were maintained on a C57BL6J background. Mice were housed under a 12h lightdark cycle and provided with food and water ad libitum. After the surgical procedure for virus injection, mice were individually housed. All animal procedures were executed in accordance with institutional guidelines, and approved by the prescribed authorities (Regierungspräsidium Freiburg).

Mice of both sexes were anesthetized and fixed in a stereotaxic frame. Adeno-associated viral vectors were injected from glass pipettes connected to a pressure ejection system into the auditory cortex. After 6-8 weeks of viral expression, mice were deeply anaesthetised with isoflurane and decapitated into carbonated, ice-cold slicing solution. A vibratome was used to obtain 350 *μ*m thick coronal slices from the auditory cortex.

Slices were held in a recording chamber and perfused with ACSF. Cells were visualized for patching using differential interference contrast microscopy or under epifluorescence for identification using an LED with a water immersion objective and a CCD camera. Cells were recorded in whole-cell patch clamp recordings using pipettes pulled from standard-wall borosilicate capillaries using a universal electrode puller. A Multiclamp 700B amplifier was used for whole-cell voltage-clamp recordings, together with a Digidata1550 for digitization. To study presynaptic GABA_B_ receptor-mediated inhibition while blocking putative postsynaptic effects of GABA_B_ receptor activation, L1 INs were recorded with Cesium-based intracellular solution. In these experiments, cells were recorded at 0 mV in control conditions, after application of baclofen and after addition of CGP55845. SOM IN inputs in L1 were optically stimulated with either 2 pulses of 0.5 ms at 10 or 20 Hz, or a naturalistic train of 10 pulses of 0.5 ms mimicking activity recorded from a L1 IN in vivo.

For microscopic analysis, the brain slices were incubated overnight following the acquisition. Slices were stained with DAPI, and mounted on objective slides before being imaged with a microscope.

Detailed descriptions of the experimental procedure can be found in the Supplementary Information.

## Data and code availability

The data and code for models and data analysis will be made available upon publication of the manuscript.

## Acknowledgments

We thank all members of the Letzkus lab, the Sprekeler lab and the Vogels lab for discussions, U. Thirimanna for technical assistance, and K. Deisseroth for generously sharing reagents. This work was supported by the German Research Foundation (LE 3804/3-1, LE 3804/4-1, LE 3804/7-1, and TRR 384/1 2024, – 514483642) and the Wellcome Trust Senior Research Fellowship 214316/Z/18/Z.

## Supplementary Information

### Supplementary Methods: Model and Simulations

#### Cortical microcircuit model with NDNF INs

Our cortical microcircuit model is a rectified, linear rate-based network consisting of excitatory PCs (*N*_E_ = 70) and the four IN types NDNF, SOM, VIP and PV (*N*_N_ = *N*_S_ = *N*_V_ = *N*_P_ = 10). All neurons are randomly connected with connections strengths and connection probabilities that are consistent with electrophysiological recordings (see ‘Connectivity’).

Excitatory PCs consist of two compartments representing the dendrite and the soma, whereas inhibitory neurons are point neurons described by a single activity value. Neural activities in our model are unitless but could be interpreted as firing rates. Note that their exact magnitudes can be arbitrarily rescaled through parameters in the network.

The activity vectors of the INs (**r**_N_, **r**_S_, **r**_V_ and **r**_P_) and the somatic and dendritic activity vectors of the PCs (**r**_E_ and **v**_D_) evolve according to

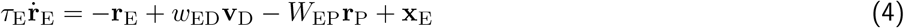

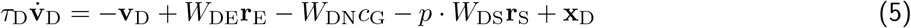

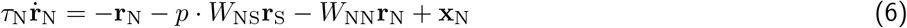

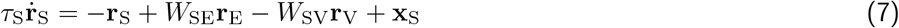

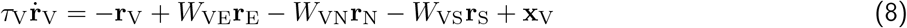

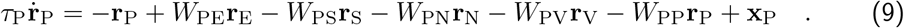

where *τ*_*X*_ are the time constants, **x**_*X*_ are external inputs strengths and *W*_*XY*_ are the synaptic weight matrices determining the connection strengths between neurons (*X, Y* ∈ {N, S, V, P, E, D} stand for NDNF, SOM, VIP, PV, PC soma and PC dendrite, respectively).

*w*_ED_ describes the coupling strength of the dendrite to the soma of the same PC and is set to 1. In addition to the linear differential equations, all activities are rectified at zero to ensure non-negative rates. See Tables 1 and 2 for an overview of the microcircuit parameters.

**Table 1.** Neuron numbers and timescales in the microcircuit model.

| population | symbol | size | symbol | timescale |
| --- | --- | --- | --- | --- |
| excitatory neurons | $N_E$ | 70 | $\tau_E$ | 10 ms |
| dendrites | - | - | $\tau_D$ | 20 ms |
| NDNF INs | $N_N$ | 10 | $\tau_N$ | 40 ms |
| SOM INs | $N_S$ | 10 | $\tau_S$ | 20 ms |
| VIP INs | $N_V$ | 10 | $\tau_V$ | 15 ms |
| PV INs | $N_P$ | 10 | $\tau_P$ | 10 ms |
| GABA concentration | - | - | $\tau_G$ | 200 ms |
| release factor | - | - | $\tau_{pi}$ | 100 ms |

**Table 2.** Default parameters of the microcircuit model.

| parameter | symbol | value |
| --- | --- | --- |
| dendrite-soma coupling | $w_{ED}$ | 1 |
| GABA release strength | $\gamma$ | 1 |
| presynaptic inhibition strength | $b$ | 0.5 |
| synaptic weight heterogeneity | $\sigma_w$ | 0.1 |
| background noise mean | $\mu_s$ | 0 |
| background noise variability | $\sigma_s$ | 0.1 |

The variables *c*_G_ and *p* relate to GABAergic volume transmission and presynaptic inhibition, respectively, and will be elaborated on in the following section.

### GABAergic volume transmission and presynaptic inhibtion

To model the effect of NDNF IN-mediated GABAergic volume transmission, we introduce the ambient GABA concentration *c*_G_. We assume that the microcircuit we consider is sufficiently small and the GABAergic volume release sufficiently broad such that the GABA concentration can be described by a single variable. The GABA concentration increases with the activity of all NDNF INs in the circuit:

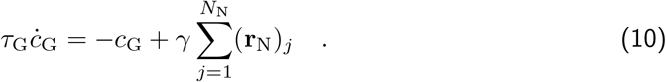

The amount of GABA released by NDNF INs (i.e., how much their activity contributes to the GABA concentration) can be scaled by the parameter *γ*, though in practice we set *γ* = 1. To account for the slower dynamics of GABAergic volume release compared to synaptic transmission, the time constant of the GABA concentration *τ*_G_ is set to 200ms, which is larger than the neural time constants (cf. Table 1). In line with experimental observations, the inhibition of NDNF INs onto PC dendrites in Eq. (5) is mediated by GABAergic volume release as described by *c*_G_.

We introduce presynaptic inhibition by decreasing the release factor *p* with increasing GABA concentration, thereby capturing the inhibition of synaptic transmission via presynaptic GABA_B_. *p* evolves according to

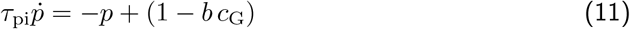

and is clipped between 0 and 1. The parameter *b* describes the strength of presynaptic inhibition (i.e. how much *p* decreases with the GABA concentration) and is set to 0.5. The timescale *τ*_p_ is set to 100ms to capture the slow dynamics of GABA_B_ receptors.

We scale the influence of SOM INs on PC dendrites and NDNF INs by the release factor *p* (see Eqs. (5)–(6)) such that the output synapses of SOM INs are modulated by presynaptic inhibition, but not the activity of SOM INs or their outputs in lower layers (e.g. SOM-to-PV inhibition).

### Connectivity

All neurons are randomly connected with connection probabilities and connection strengths motivated by the experimental literature (Pfeffer et al., 2013; Pi et al., 2013; Jiang et al., 2015; Abs et al., 2018; Campagnola et al., 2022; Hartung et al., 2023). The matrix of connection probabilities is given by

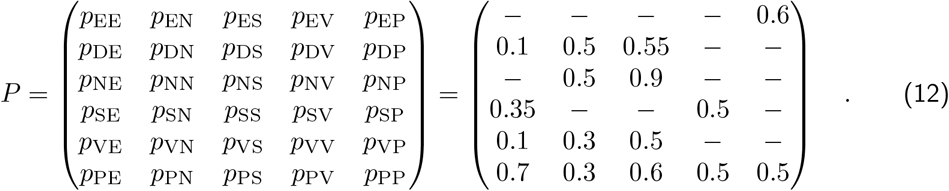

All cells of the same neuron type have the same number of incoming connections, that is, the in-degree for each connection type is fixed. Connectivity is random, except that neurons do not connect to themselves (i.e. no autapses). The default mean synaptic connection strengths are given by

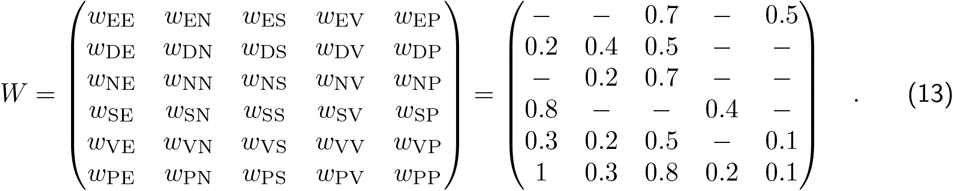

All weights are scaled by the number of incoming connections for each connection type such that the results are independent of the population size. This means that the effective mean weight is 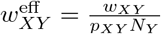, where *N*_*Y*_ is the presynaptic population size. In addition, we scale the connections affected by presynaptic inhibition by the release factor at baseline *p*_0_ such that the baseline steady-state activities are independent of the presynaptic inhibition strength. This means that the mean SOM-to-NDNF and SOM-to-dendrite synaptic weights are 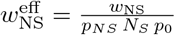 and 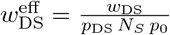 where *p*_0_ = 1 − *b* **r**_N_ = 0.5 for the default network parameters.

Individual weights in the weight matrices *W*_*XY*_ (as used in Eqs. (4)–(9)) are sampled from Gaussian distributions 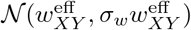 with *σ*_*w*_ = 0.1. We ensure that all weights are positive by rectifying the randomly sampled weight values.

### Inputs

All neurons receive constant, external background input that ensures non-zero baseline activity. We tune the background inputs **x**_*X*_ in Eqs. (4)–(9) such that all neurons have an average baseline activity of 1. To this end, we consider a mean-field version of our model, in which each population is represented by a single activity variable. Since we use a rate model, this is equivalent to setting the number of neurons per type to 1 and removing the heterogeneity in synaptic weights (i.e. *σ*_*w*_ = 0). We then compute the steady-state for each population, insert the desired target activities, and solve the equations for *x*_E_, *?*_D_, *x*_N_, *x*_S_, *x*_V_ and *x*_P_. Note that in the mean-field model the background inputs *x*_*X*_ are scalars, whereas in the full microcircuit model, the inputs **x**_*X*_ are vectors with **x**_*X*_ = *x*_*X*_ **1**. The default background input parameters are listed in Table 3. The inputs are unitless, consistent with the unitless neural activities, but could be interpreted as firing rates, for instance. Not that the exact values need to be re-computed for changes in mean synaptic weights or other network parameters.

**Table 3.** Default background inputs.

| population | symbol | input strength |
| --- | --- | --- |
| excitatory neurons | $x_E$ | 0.5 |
| dendrites | $x_D$ | 1.7 |
| NDNF INs | $x_N$ | 1.9 |
| SOM INs | $x_S$ | 0.6 |
| VIP INs | $x_V$ | 1.4 |
| PV INs | $x_P$ | 1.4 |

To account for variability in external inputs we add time-varying Gaussian white noise with mean *μ*_*s*_ = 0 and standard deviation *σ*_*s*_ = 0.1 to the background inputs.

### Manipulations of the microcircuit

Above we described the default microcircuit model and its inputs. To study the functional implications of controlling SOM outputs via GABAergic volume transmission, we perform a range of manipulations of the circuit. Below we provide a detailed description of the *in silico* protocols for each figure containing manipulations of the default circuit model.

**Figure 3: Competition between SOM-and NDNF-mediated dendritic inhibition**. We varied the input to NDNF INs relative to their background input. This was achieved by providing an additional positive or negative input to NDNF INs (Δ NDNF input, Fig. 3). To change the relative strength of SOM-to-dendrite compared to NDNF-to-dendrite inhibition, we varied the mean unscaled NDNF-to-dendrite synaptic weight *w*_DN_ between 0 and 0.8. To remove presynaptic inhibition from the model, we set *b* = 0 such that *p* = 1 at all times (Eq. (11)). Amplification was tested for different levels of SOM-to-NDNF inhibition *w*_NS_ between 0.5 and 1.7. We quantified the amplification of inputs to NDNF INs by presynaptic inhibition using an amplification index (Hertäg and Sprekeler, 2019). To this end, we fitted lines to the linear parts of the curves in Fig. 3C (Δ NDNF input between − 0.3 and 0.3) with and without presynaptic inhibition. The amplification index is then given by the logarithmic ratio of the two slopes:

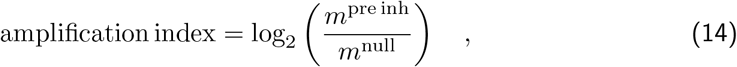

where *m*^preinh^ and *m*^null^ are the slopes of the linear fit with and without presynaptic inhibition, respectively.

**Figure 4: NDNF INs can act as a switch for dendritic inhibition**. We provided a positive pulse input of 0.6 or a negative pulse input of − 0.5 for 1 second to NDNF INs to switch between NDNF-and SOM-mediated dendritic inhibition. We then measured the steady-state NDNF activity five seconds after the pulse. In Fig. 4C we varied the strength of this pulse between −1 and 1. To ensure that PC activity does not change when dendritic inhibition is switched between NDNF-and SOM-mediate, we increased the NDNF-to-dendrite inhibition by setting *w*_DN_ = 0.6 (default is 0.4, see Eq. (13)). For weak SOM-to-NDNF inhibition, we used the default value *w*_NS_ = 0.7, and for strong SOM-to-NDNF inhibition, we used *w*_NS_ = 1.2. Note that changing the mean synaptic weights requires recomputing the background inputs to maintain the same baseline neural activities. In Fig. 4D we varied the strength of this inhibition between 0.5 and 1.6. The time-varying stimulus provided to SOM INs in Fig. 4F was a sine wave with frequency 2Hz and amplitude 0.5.

**Figure 5: Redistribution of dendritic inhibition in time**. To demonstrate the different temporal scales of SOM and NDNF inhibition we provided a brief stimulus to each cell type and measured the elicited postsynaptic current in PCs, mimicking a paired recording protocol *in silico* (Fig. 5B). The stimulus was an instantaneous increase followed by an exponential decay at 50 ms. Next, we tested the frequency response curve of PCs upon stimulating SOM and NDNF INs with sine waves of different frequencies (Fig. 5C) at an amplitude of 1.5. We measured the signal transmission to PCs by computing the amplitude of the elicited oscillation (mean of peak-to-valley response). Finally, we tested how stimulating SOM and NDNF INs with constant stimuli of different durations affects PCs (Fig. 5D). Both cell types were stimulated with a constant additive input of 1.5 that lasted for intervals ranging from 10ms to 1s. We measured the response in PCs by averaging over the time window of the stimulus. The change in different inhibitory pathways was obtained by recording the respective currents into PCs (weight times activity) and computing the average difference before and during the stimulus. We repeated this experiment for weak (i.e. default) and strong NDNF-to-dendrite inhibition (*w*_DN_ = 0.4 and *w*_DN_ = 0.8, respectively). To remove presynaptic inhibition from the model, we set *b* = 0.

### Model changes for supplementary figures

We tested the robustness of our results to variations of the microcircuit architecture and parameters. A summary of the model changes in the supplementary figures can be found in Table 4.

**Table 4.** Overview of model changes in supplementary figures.

| Figure | Result | Model change |
| --- | --- | --- |
| Supp. Fig. 2 | The results are robust to presynaptic inhibition targeting NDNF-dendrite synapses. | NDNF-to-dendrite weight $w_{\text{DN}}$ is multiplicatively modulated by $p^{\text{DN}}$, which changes similar to $p$ in Eq. (11) but with $b = 0.25$ |
| Supp. Fig. 3 | The results are robust to presynaptic inhibition targeting SOM-VIP synapses. | SOM-to-VIP weight $w_{\text{VS}}$ is multiplicatively modulated by $p$ from Eq. (11) |
| Supp. Fig. 4 | Temporal responses of the circuit to NDNF stimulation for weak NDNF-dendrite inhibition. | No model change, $w_{\text{DN}} = 0.4$ is the default in Eq. 13. |
| Supp. Fig. 5 | Temporal responses of the circuit to NDNF stimulation for strong NDNF-dendrite inhibition. | NDNF-to-dendrite weight increased to $w_{\text{DN}} = 0.8$. |
| Supp. Fig. 7 | Without presynaptic inhibition activating NDNF INs does not qualitatively change mismatch responses of PCs. | No presynaptic inhibition, i.e. $b = 0$ and $p = 1$. |
| Supp. Fig. 8 | The modulation of the prediction error circuit through NDNF activation is robust to model variations. | A: NDNF-to-PV weight $w_{\text{PN}}$ is modulated by $p$. B: NDNF INs receive $I^{\text{P}}$ as additional input. |

### Predictive coding microcircuit

For the predictive coding microcircuit (Fig. 6), we tuned the model parameters such that excitatory neurons (PCs) are mismatch neurons (Hertäg and Sprekeler, 2020). The circuit received sensory input mimicking, e.g. visual flow, and prediction input representing the expected sensory input based on, e.g. motor commands (Attinger et al., 2017; Keller and Mrsic-Flogel, 2018; Hertäg and Sprekeler, 2020). Sensory input *I*^S^ targeted the PC soma, SOM and PV INs, whereas the prediction *I*^P^ targeted PC dendrites and VIP INs (Fig. 6). Since NDNF INs are targeted by a range of top-down projections, they may also receive prediction input. For simplicity we here assume that this is not the case, however, our results still hold when NDNF INs also receive the prediction input *I*^P^ (Supp. Fig. S8B). The equations describing the dynamics of the predictive coding circuit are the same as in Eqs. (4)–(11), just with the additional inputs *I*^S^ and *I*^P^ to the respective cell types. The amplitude of the prediction and the sensory input was 1 when the respective input was present and 0 otherwise (cf. Fig. 6).

PCs are considered mismatch neurons if they only respond when the prediction is larger than the sensory input (*I*^P^ *> I*^S^, “mismatch”) but not when the sensory input is larger (*I*^P^ *< I*^S^, “playback”) or when the sensory input matches the prediction (*I*^P^ = *I*^S^, “feedback”). This can be achieved by balancing multiple inhibitory, disinhibitory and excitatory pathways in the microcircuit (Hertäg and Sprekeler, 2020). We here used the mean weight parameters derived by Hertäg and Sprekeler (2020) as a starting point. However, the original predictive coding circuit did not contain NDNF INs. We included NDNF INs with the same weight parameters as for our default microcircuit model and then hand-tuned their strengths to ensure that PCs produce prediction error neuron responses. The adapted mean synaptic weight parameters are given by

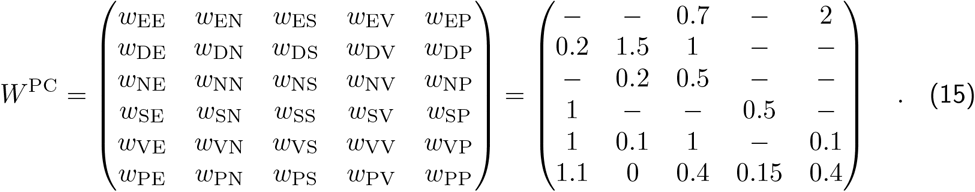

Connection probabilities were the same as before. For simplicity, we assumed that the connection from NDNF to PV INs is absent in the predictive coding circuit (*w*_PN_ = 0). This makes balancing the multiple pathways easier because, without NDNF-to-PV inhibition, changes in SOM IN activity only affect PV INs directly and not indirectly through the disinhibitory SOM-NDNF-PV pathway. Indeed, NDNF-to-PV inhibition is substantially weaker than NDNF-to-dendrite inhibition (Hartung et al., 2023), albeit non-zero. We show that PCs can still be prediction error neurons, even when *w*_PN_ *>* 0 (Supp. Fig. S8A).

For consistency with the predictive coding microcircuit by Hertäg and Sprekeler (2020), we tuned the background inputs such that at baseline all IN activities are 4, the PC activity is 1 and the dendrite is inactive (activity is 0). The corresponding mean background inputs are *x*_E_ = 9, *x*_D_ = 9.8, *x*_N_ = 6.8, *x*_S_ = 5, *x*_V_ = 7.4 and *x*_P_ = 6.2. To accommodate for the increased baseline activity of NDNF INs, we reduced the strength of presynaptic inhibition to *b*^PC^ = 0.15 such that the release factor is *p*_0_ = 0.4 in the absence of predictions or sensory input. Note that as in the default microcircuit model, connections affected by presynaptic inhibition need to be scaled by population size, connection probability, and initial release probability *p*_0_ (see ‘Connections’).

To mimic elevated NDNF IN activity in response to neuromodulatory input from, e.g. cholinergic projections, we provide an additional additive input to NDNF INs (Brombas et al., 2014; Poorthuis et al., 2018). The magnitude of this additional input is 1.

### Simulation details

All simulations were performed in customised Python code written by LBN. Differential equations were numerically integrated using the forward Euler method with a time step of 1 ms. Neurons were initialised at their analytical steady-state (i.e. baseline target) value. The simulation duration varied between different experiments.

## Supplementary Methods: Experimental protocol

### Animal subjects

All mouse lines used were maintained on a C57BL6/J background. Mice were housed under a 12h light/dark cycle and provided with food and water ad libitum. After the surgical procedure for virus injection, mice were individually housed. All animal procedures were executed in accordance with institutional guidelines, and approved by the prescribed authorities (Regierungspräsidium Freiburg).

### Surgery

Mice were anaesthetised with isoflurane (induction: 5%, maintenance: 1.5-2%) in oxygen-enriched air (Oxymat 3, Weinmann, Hamburg, Germany) and fixed in a stereotaxic frame (Kopf Instruments, Tujunga, USA). Core body temperature was maintained around 37-38^*°*^C via a feed-back controlled heating pad (FHC, Bowdoinham, ME, USA). Analgesia was provided by local injection of ropivacain under the scalp (16.7mg/kg, Ropivacain-HCl B.Braun) and subcutaneous injection for systemic action of metamizol (200 mg/kg, Novaminsulfon-ratiopharm) and meloxicam (1-2 mg/kg, Metacam Boehringer Ingelheim). Adeno-associated viral vectors (AAV2/5.EF1a.DIO.hChR2(H134R)-EYFP.WPRE.hGH, PennVectorCore; pAAV-Ef1a-fDIO-tdTomato (AAV1), Addgene; 1:2 mix, 600 nl injected) were injected from glass pipettes connected to a pressure ejection system (PDES-02DELA-2, NPI, Germany) into auditory cortex at the following coordinates: 2.54 mm posterior from bregma, 4.6 mm lateral of midline, depth below cortical surface 100-900 *μ*m.

### Slice preparation

Mice of both sexes (12-16 weeks, 6-8 weeks viral expression) were deeply anaesthetised with isoflurane (5%) in oxygen-enriched air (Oxymat 3, Weinmann, Hamburg, Germany), and decapitated into carbonated, ice-cold slicing solution. A Leica VT 1200S vibratome was used to obtain 350 *μ*m thick coronal slices from auditory cortex. Slices were directly transferred to carbonated slicing solution at 33^*°*^C for 15 minutes and then transferred to carbonated standard ACSF at room temperature. After 30-60 minutes of recovery time, slices were used in whole-cell patch-clamp experiments. Slicing solution contained (in mM) 93 NMDG, 93 HCl, 2.5 KCl, 1.2 NaH2PO4, 30 NaHCO3, 20 HEPES, 25 glucose, 5 sodium ascorbate, 2 thiourea, 3 sodium pyruvate, 10 MgSO4 and 0.5 CaCl2 and was calibrated to a pH of 7.3-7.4 and an osmolarity of 300-310 mOsm. Standard ACSF contained (in mM) 125 NaCl, 3 KCl, 1.25 NaH2PO4, 26 NaHCO3, 10 glucose, 1 MgCl2 and 2 CaCl2 and was calibrated to an osmolarity of 300-310 mOsm.

### Patch clamp electrophysiology

Slices were held in a recording chamber at 33^*°*^C and perfused with ACSF (2-4 mL/min). Cells were visualized for patching using differential interference contrast microscopy (Scientifica) or under epifluorescence for identification using an LED (488 or 565 nm, Cool LED) with a water immersion objective (40x, 0.8 N.A., Olympus LUMPLFLN) and a CCD camera (Hamamatsu C11440 ORCA-flash4.0). Cells were recorded in whole-cell patch clamp recordings using pipettes pulled from standard-wall borosilicate capillaries using a universal electrode puller (3.5-6 MOhm, DMZ Zeitz-Puller). A Multiclamp 700B amplifier (Axon Instruments, CA) was used for whole cell voltage clamp recordings, together with a Digidata1550 (Molecular Devices) for digitization. Recordings were low pass filtered at 10 or 2.4 kHz using a Bessel filter and digitized at 20 kHz. Series resistance was routinely compensated in voltage clamp and recordings were excluded when access resistance exceeded 50 MOhm. To study presynaptic GABA_R_ receptor-mediated inhibition while blocking putative postsynaptic effects of GABA_R_ receptor activation, L1 INs were recorded with Cesium-based intracellular solution containing (in mM): 130 CsOH, 130 D-gluconic acid, 2 MgCl2, 0.2 EGTA, 5 NaCl, 10 HEPES, 4 ATP-tris, 0.3 GTP-tris, 10 phosphocreatine (pH 7.3, 290 mOsm). In these experiments, cells were recorded at 0 mV in control conditions, after application of baclofen (10 *μ*M, Tocris) and after addition of CGP55845 (3 *μ*M, Sigma). SOM IN inputs in L1 were optically stimulated (488 nm) with either 2 pulses of 0.5 ms at 10 or 20 Hz, or a naturalistic train of 10 pulses of 0.5 ms mimicking activity recorded from a L1 IN in vivo (data not shown), containing instantaneous frequencies between 1 and 26.77 Hz.

### Microscope analysis of brain slices

Slices were incubated in 4% PFA overnight at 4^*°*^C following the acquisition. Following three wash steps of 10 minutes with PBS, slices were stained with DAPI (0.5 mg/ml, D1306, Thermo Fisher Scientific) for 10 minutes, after which the same wash procedure followed. Finally, slices were mounted on objective slides and mounted with Mowiol 4-88 (Polysciences) before being imaged with a Zeiss confocal microscope (Axio Zoom, LSM 790).

### Statistical analysis

Statistical analysis was performed using GraphPad Prism. Data were considered normally distributed if Shapiro-Wilk, D’Agostino & Pearson and KS tests were passed. According to this result, and depending on whether data was paired or not, comparisons were performed using the following parametric or non-parametric tests: For three-group comparisons, One-way ANOVA (normal, non-paired) or One-way repeated measures (RM) ANOVA (normal, paired) followed by Tukey’s multiple comparisons test, and Kruskal-Wallis test (non-normal, non-paired) or Friedman test (non-normal, paired) followed by Dunn’s multiple comparisons test. Two-way RM ANOVA followed by Sidak’s and Tukey’s multiple comparison tests were used to compare groups across more than one factor. Statistical tests used in each instance are indicated in the figure legends and results are reported as follows: n.s. (not significant) *p >* 0.05, * *p <* 0.05, ** *p <* 0.01, *** *p <* 0.001, **** *p <* 0.0001.

## Supplementary Figures

**Figure S1.**
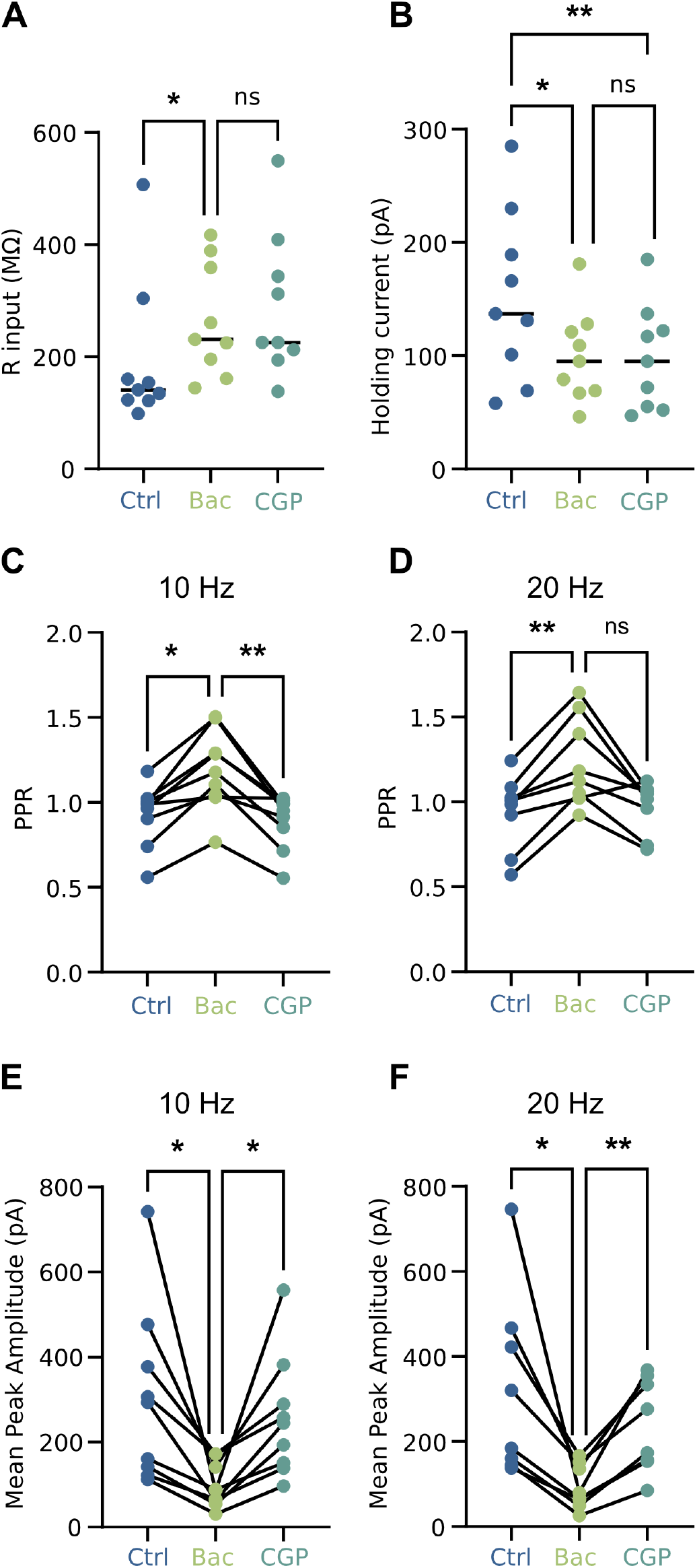
**A**. Input resistance and B. holding current recorded for control (ACSF), baclofen and CGP55845 conditions. As the input resistance and holding current do not change between baclofen and CGP55845, the effect on IPSC amplitude and PPR shown in Fig. 2 cannot result from postsynaptic effects. **C/D**. PPR recorded for control (ACSF), baclofen and CGP55845 conditions with 10 Hz and 20 Hz, respectively. **E/F**. Mean peak amplitude recorded for control (ACSF), baclofen and CGP55845 conditions with 10 Hz and 20 Hz, respectively. Data shown as averages of 10 sweeps and median (A,B) or without median (C, D, E, F). n.s. *P >* 0.05, * *P <* 0.05, ** *P <* 0.01, *** *P <* 0.001.

**Figure S2.**
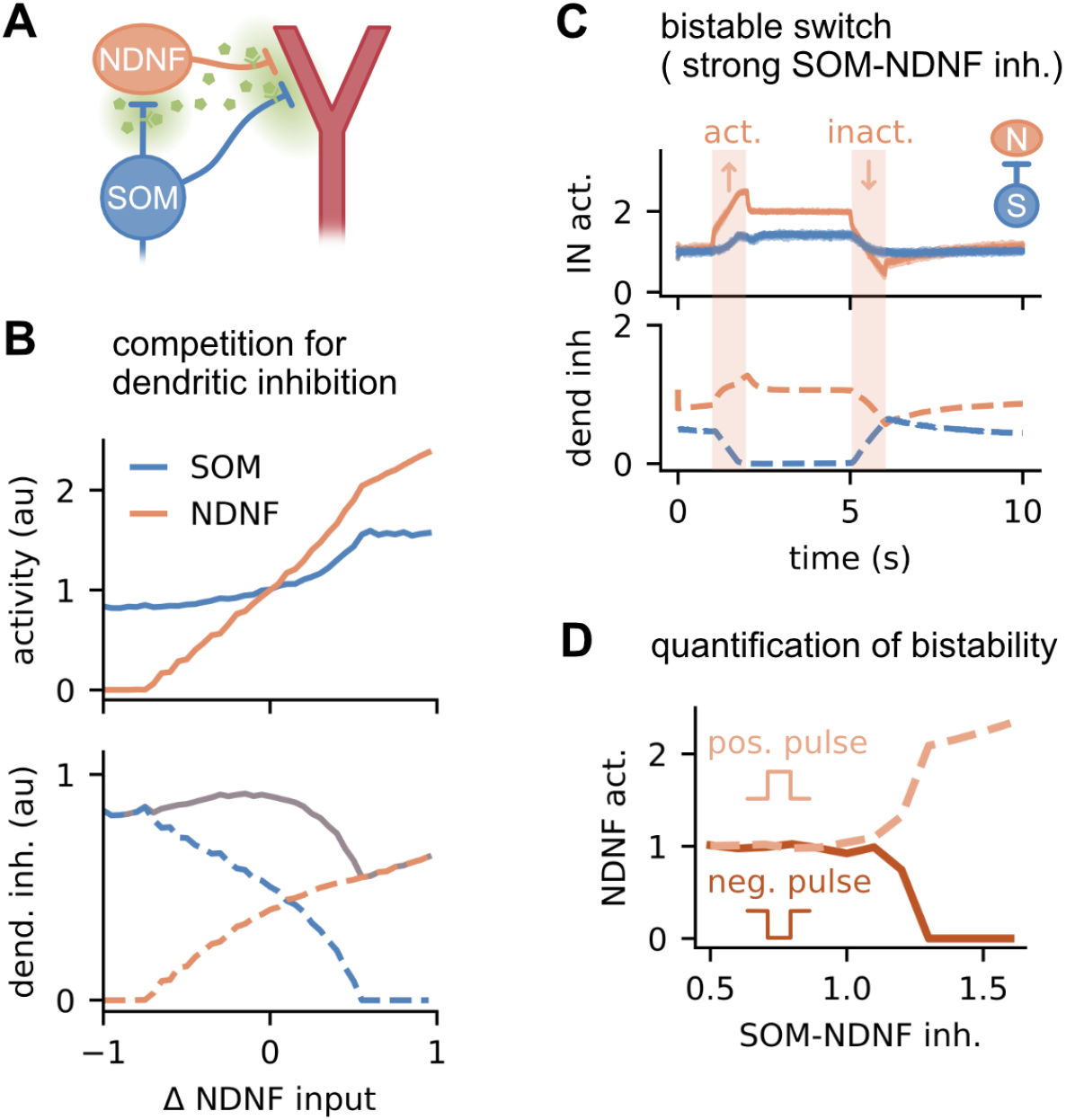
The results are robust to presynaptic inhibition targeting NDNF-dendrite synapses. **A**.Schematic of the model with presynaptic inhibition targeting NDNF-to-dendrite synapses. **B**.Activity of NDNF and SOM INs (top) and dendritic inhibition from SOMs, NDNFs and both combined (grey; bottom) as a function of NDNF input. Colours correspond to the IN colours in (A). **C**. Time course of SOM IN and NDNF IN activity (top) and the dendritic inhibition they exert (bottom) when NDNF INs are switched on and off. SOM-NDNF inhibition is strong (*w*_NS_ = 1.2). **D**. Steady-state NDNF IN activity as a function of SOM-NDNF inhibition strength after a positive (dashed) or negative (solid) pulse to NDNF INs.

**Figure S3.**
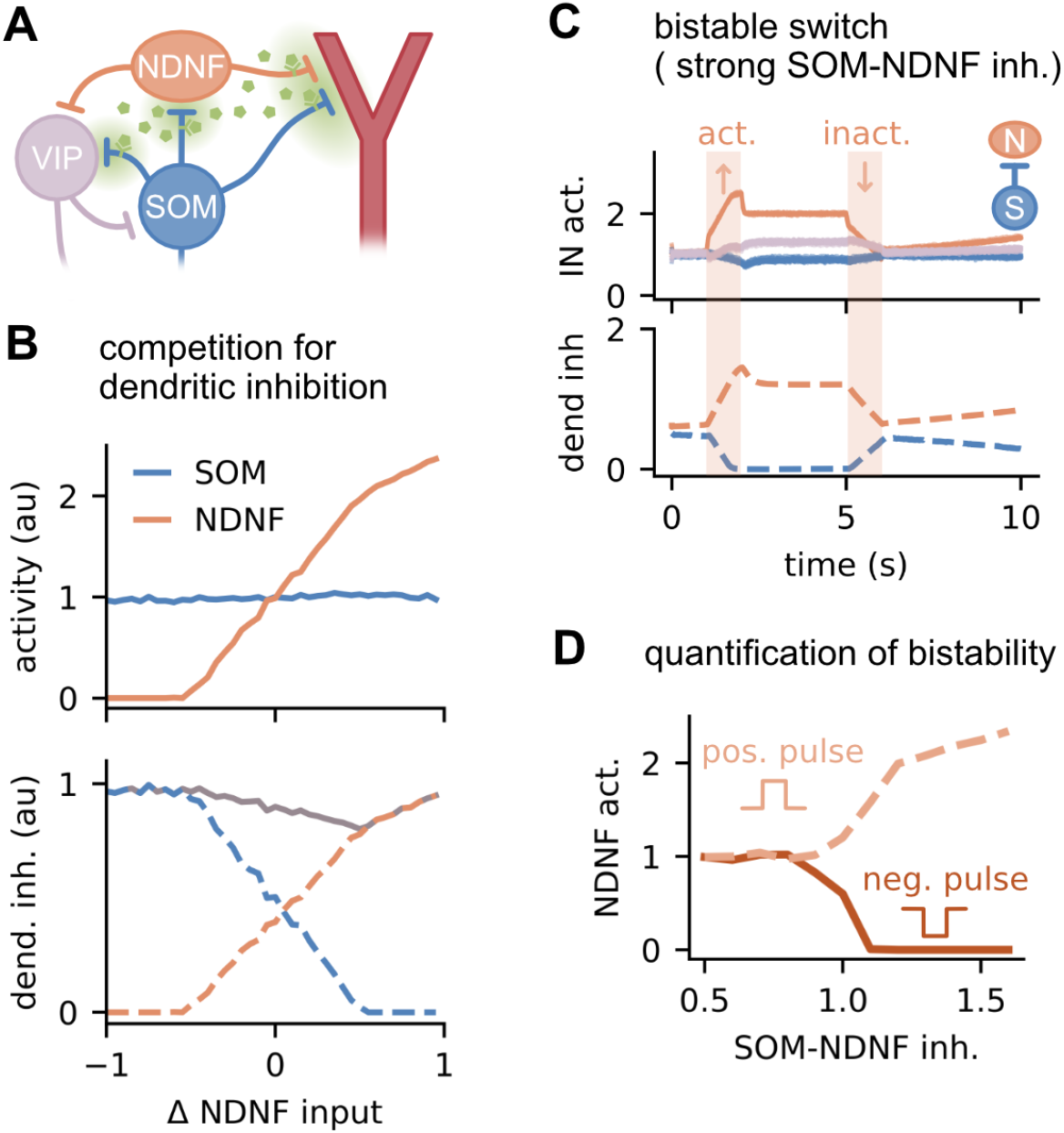
The results are robust to presynaptic inhibition targeting SOM-VIP synapses. **A**. Schematic of the model with presynaptic inhibition targeting SOM-VIP synapses. **B**. Activity of NDNF and SOM INs (top) and dendritic inhibition from SOMs, NDNFs and both combined (grey; bottom) as a function of NDNF IN input. Colours correspond to the IN colours in (A). Time course of SOM IN and NDNF IN activity (top) and the dendritic inhibition they exert (bottom) when NDNF INs are switched on and off. SOM-NDNF inhibition is strong (*w*_NS_ = 1.2). **D**. Steady-state NDNF IN activity as a function of SOM-NDNF inhibition strength after a positive (dashed) or negative (solid) pulse to NDNF INs.

**Figure S4.**
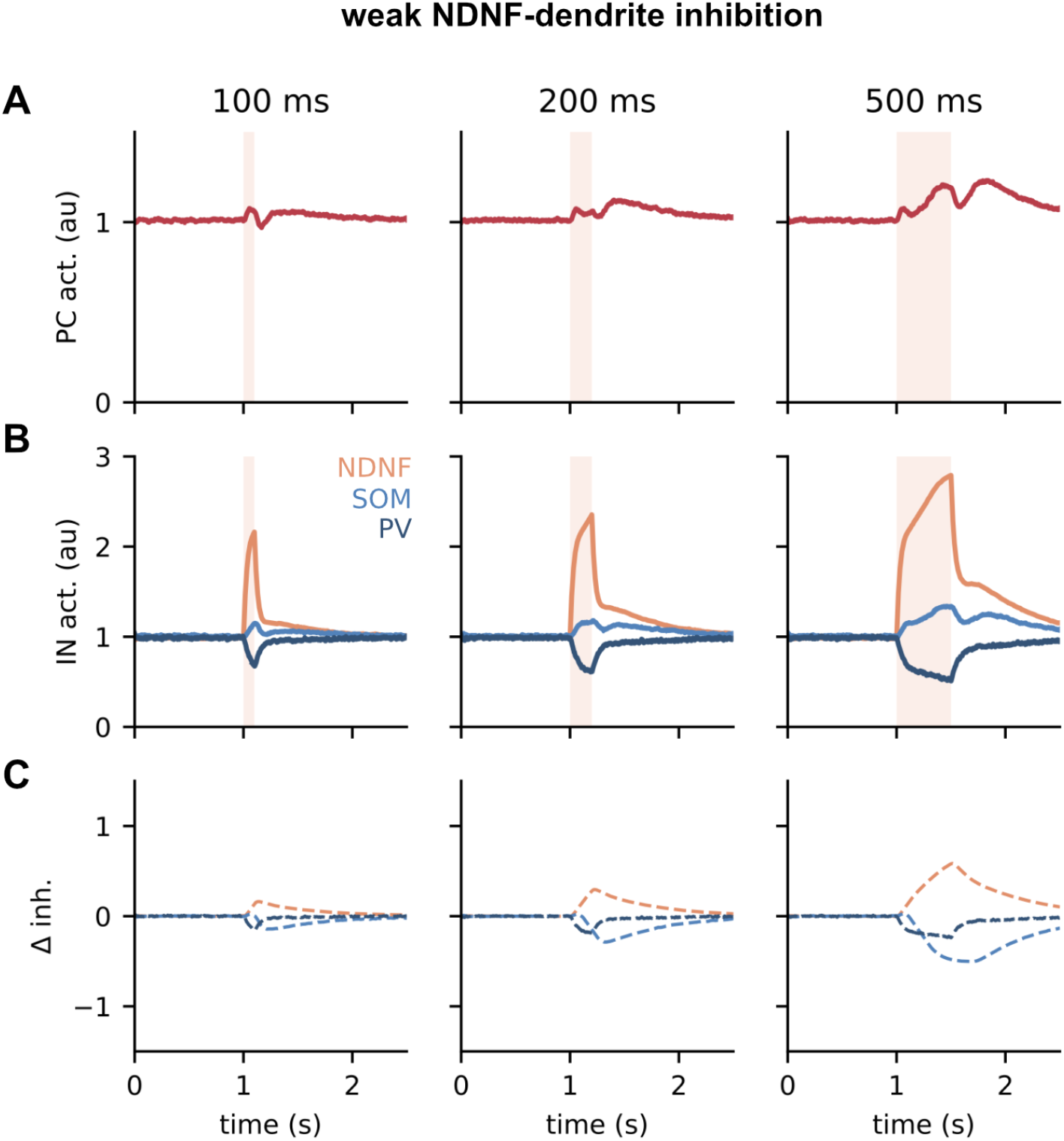
Temporal responses of the circuit to NDNF stimulation for weak NDNF-dendrite inhibition (*w*_DN_ = 0.4). **A**. PC activity. **B**. Activity of NDNF, SOM and PV INs. **C**. Change in inhibition to the PC in response to NDNF stimulation. Colours correspond to the legend in (B). Left, center and right show NDNF IN stimulation of different lengths (100, 200 and 500 ms).

**Figure S5.**
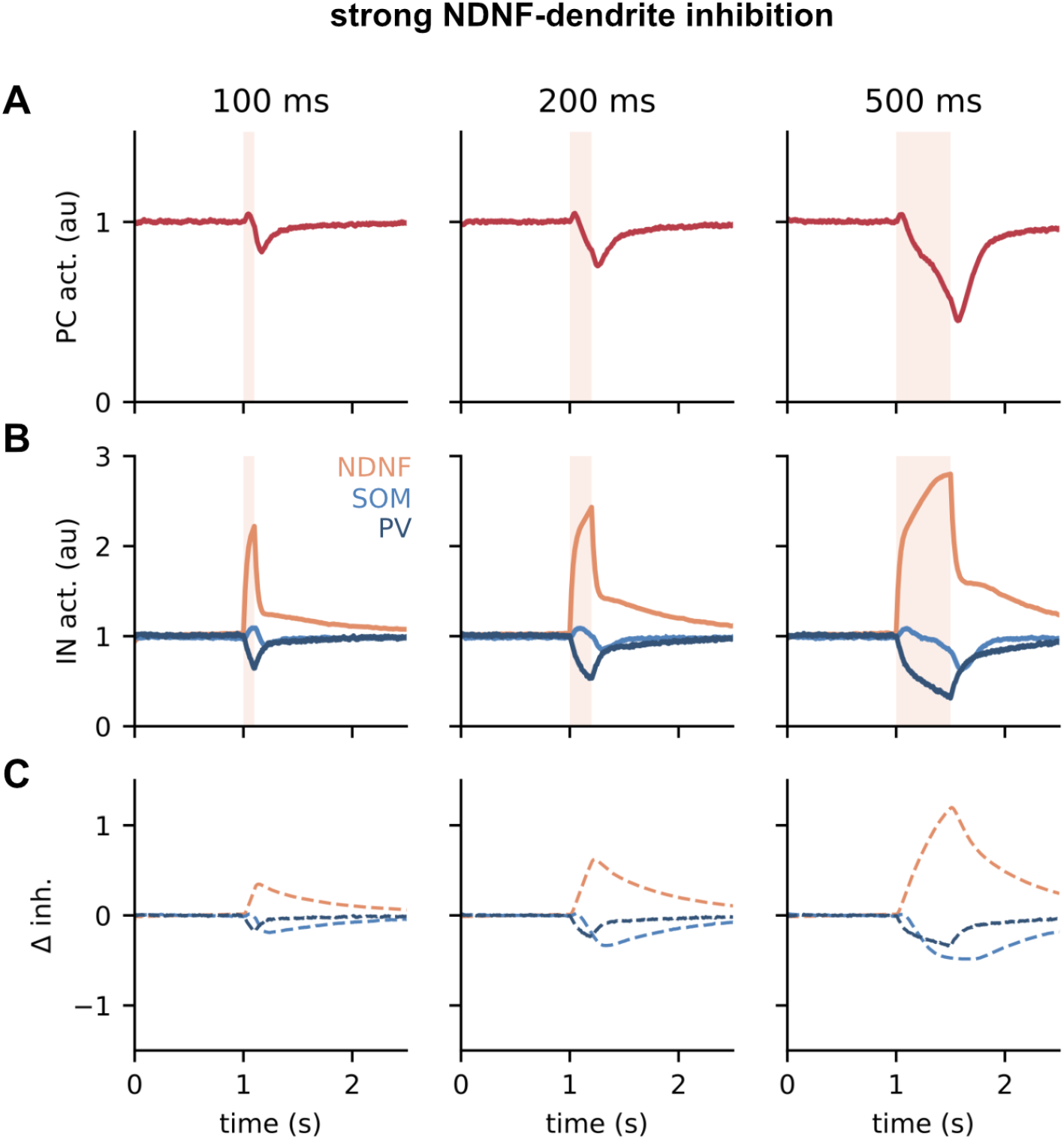
Temporal responses of the circuit to NDNF stimulation for strong NDNF-dendrite inhibition (*w*_DN_ = 0.8). **A**. PC activity. **B**. Activity of NDNF, SOM and PV INs. **C**. Change in inhibition to the PC in response to NDNF IN stimulation. Colours correspond to the legend in (B). Left, center and right show NDNF IN stimulation of different lengths (100, 200 and 500 ms).

**Figure S6.**
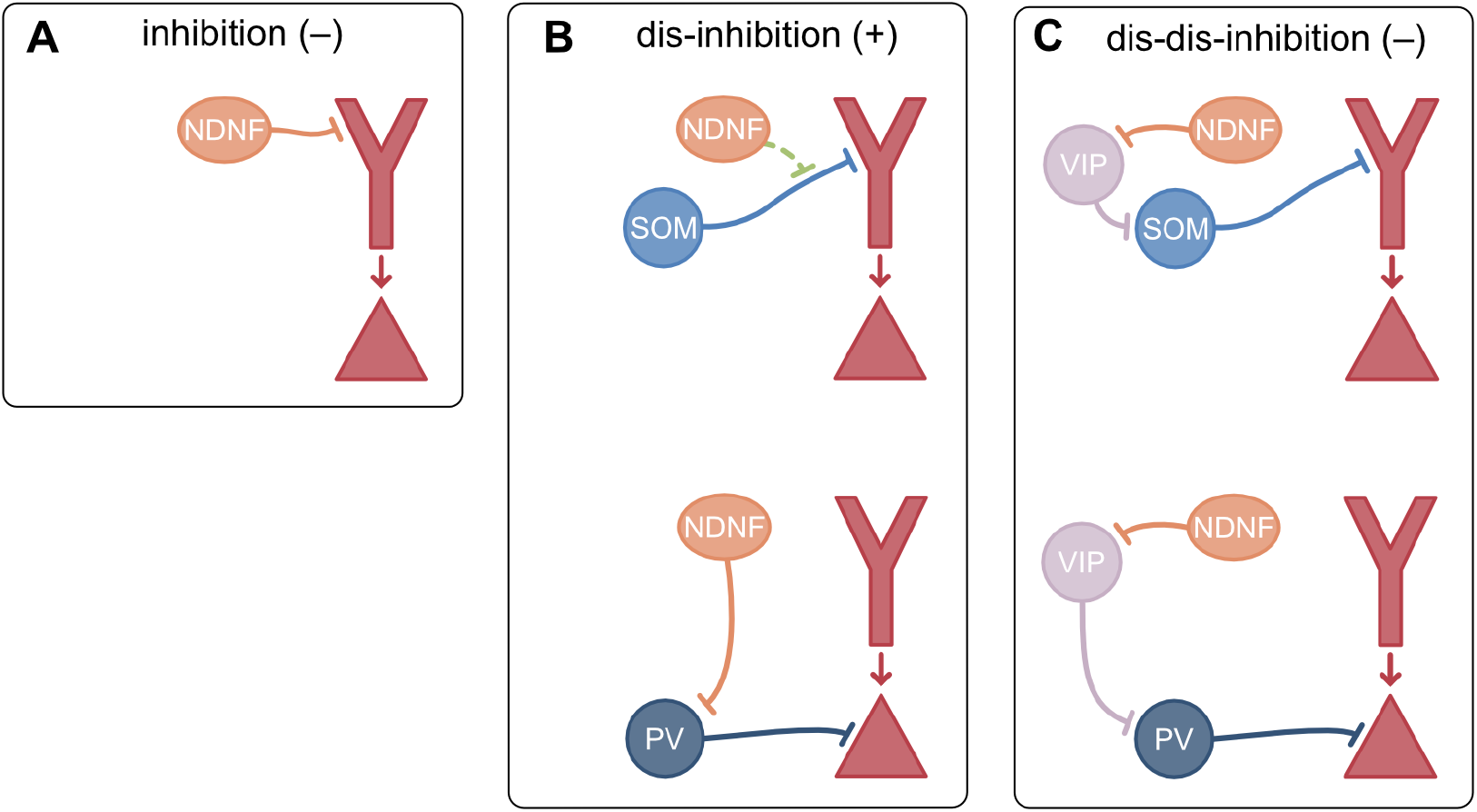
Multiple (dis-)inhibitory pathways from NDNF INs to PCs. **A**. Direction NDNF-dendrite inhibition. **B**. Presynaptic inhibition of SOM-dendrite inhibition (top) and disinhibition through the NDNF-PV-PC pathway. **C**. Dis-dis-inhibition through the NDNF-VIP-NDNF-dendrite pathway (top) and the NDNF-VIP-PV-PC pathway (bottom).

**Figure S7.**
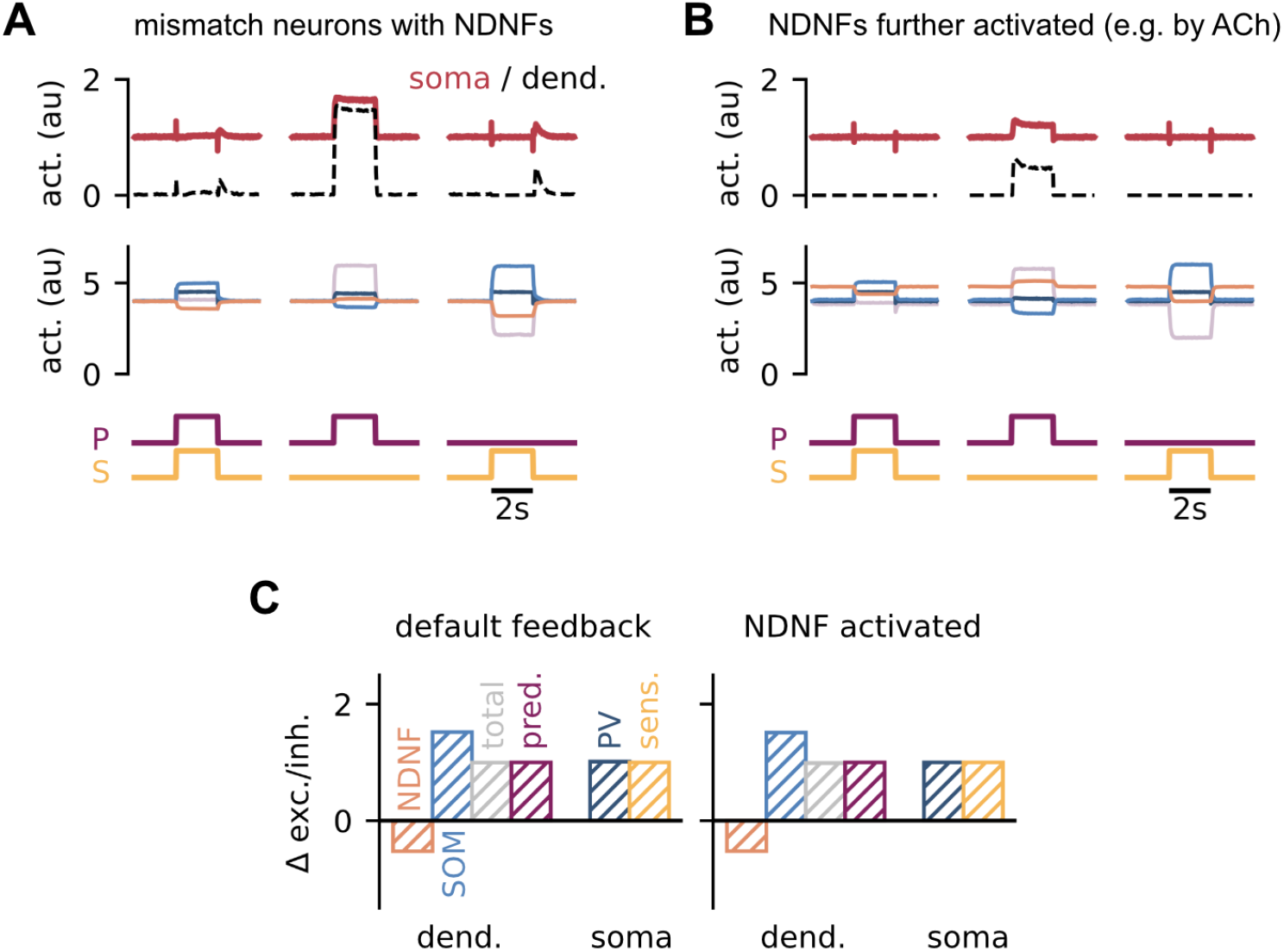
Without presynaptic inhibition activating NDNF INs does not qualitatively change mismatch responses of PCs. **A**. Responses of PC dendrites and somata (top) and four IN groups (center) to different input configurations. IN types: NDNF (orange), SOM (light blue), PV (dark blue) and VIP (lilac). **B**. Same as (B) but with NDNF INs activated, e.g. by cholinergic input. **C**. Change in excitatory and inhibitory inputs to dendrite and soma during the feedback phase in the default condition (left) and with NDNF INs activated (right).

**Figure S8.**
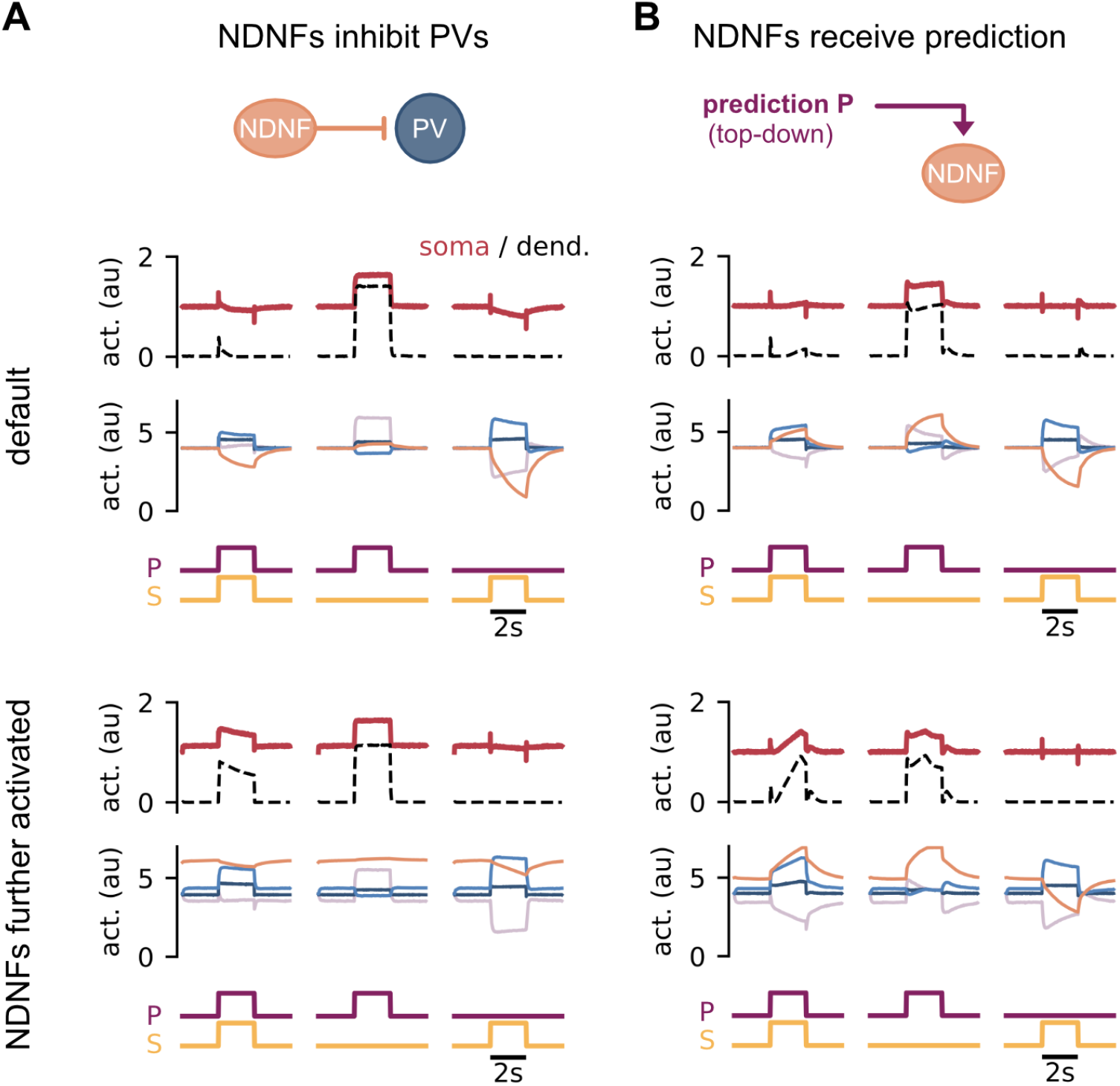
The modulation of the prediction error circuit through NDNF IN activation is robust to model variations. **A**. NDNF INs inhibit PV INs. Top: Schematic of the model change. Middle: Responses of PC dendrites, soma and the four IN groups to different input configurations in the default condition. Bottom: Same as above but with NDNF INs activated further (e.g. by cholinergic input). IN type legend: NDNF (orange), SOM (light blue), PV (dark blue) and VIP (lilac). **B**. Same as (A) but with the model change that prediction input targets NDNF INs.

